# *Wolbachia*-mediated resistance to Zika virus infection in *Aedes aegypti* is dominated by diverse transcriptional regulation and weak evolutionary pressures

**DOI:** 10.1101/2023.06.26.546271

**Authors:** Emma C. Boehm, Anna S. Jaeger, Hunter J. Ries, David Castañeda, Andrea M. Weiler, Corina C. Valencia, James Weger-Lucarelli, Gregory D. Ebel, Shelby L. O’Connor, Thomas C. Friedrich, Mostafa Zamanian, Matthew T. Aliota

**Affiliations:** Department of Veterinary and Biomedical Sciences, University of Minnesota, Twin Cities; Wisconsin National Primate Research Center, University of Wisconsin–Madison, Madison, WI, United States; Virginia Polytechnic Institute and State University, Blacksburg, VA, United States; Colorado State University, Fort Collins, CO, United States; Department of Pathology and Laboratory Medicine, University of Wisconsin–Madison, Madison, WI, United States; Department of Pathobiological Sciences, University of Wisconsin–Madison, Madison, WI, United States

**Author notes:** These authors contributed equally to this work (Emma C. Boehm, Anna S. Jaeger, Hunter J. Ries).

## Abstract

A promising candidate for arbovirus control and prevention relies on replacing arbovirus-susceptible *Aedes aegypti* populations with mosquitoes that have been colonized by the intracellular bacterium *Wolbachia* and thus have a reduced capacity to transmit arboviruses. This reduced capacity to transmit arboviruses is mediated through a phenomenon referred to as pathogen blocking. Pathogen blocking has primarily been proposed as a tool to control dengue virus (DENV) transmission, however it works against a range of viruses, including Zika virus (ZIKV). Despite years of research, the molecular mechanisms underlying pathogen blocking still need to be better understood. Here, we used RNA-seq to characterize mosquito gene transcription dynamics in *Ae. aegypti* infected with the *w*Mel strain of *Wolbachia* that are being released by the World Mosquito Program in Medellín, Colombia. Comparative analyses using ZIKV-infected, uninfected tissues, and mosquitoes without *Wolbachia* revealed that the influence of *w*Mel on mosquito gene transcription is multifactorial. Importantly, because *Wolbachia* limits, but does not completely prevent, replication of ZIKV and other viruses in coinfected mosquitoes, there is a possibility that these viruses could evolve resistance to pathogen blocking. Therefore, to understand the influence of *Wolbachia* on within-host ZIKV evolution, we characterized the genetic diversity of molecularly barcoded ZIKV virus populations replicating in *Wolbachia*-infected mosquitoes and found that within-host ZIKV evolution was subject to weak purifying selection and, unexpectedly, loose anatomical bottlenecks in the presence and absence of *Wolbachia*. Together, these findings suggest that there is no clear transcriptional profile associated with *Wolbachia*-mediated ZIKV restriction, and that there is no evidence for ZIKV escape from this restriction in our system.

**Author Summary:** When *Wolbachia* bacteria infect *Aedes aegypti* mosquitoes, they dramatically reduce the mosquitoes’ susceptibility to infection with a range of arthropod-borne viruses, including Zika virus (ZIKV). Although this pathogen-blocking effect has been widely recognized, its mechanisms remain unclear. Furthermore, because *Wolbachia* limits, but does not completely prevent, replication of ZIKV and other viruses in coinfected mosquitoes, there is a possibility that these viruses could evolve resistance to *Wolbachia*-mediated blocking. Here, we use host transcriptomics and viral genome sequencing to examine the mechanisms of ZIKV pathogen blocking by *Wolbachia* and viral evolutionary dynamics in *Ae. aegypti* mosquitoes. We find complex transcriptome patterns that do not suggest a single clear mechanism for pathogen blocking. We also find no evidence that *Wolbachia* exerts detectable selective pressures on ZIKV in coinfected mosquitoes. Together our data suggest that it may be difficult for ZIKV to evolve Wolbachia resistance, perhaps due to the complexity of the pathogen blockade mechanism.

## Introduction

The intracellular bacterium *Wolbachia pipientis* naturally infects a range of insect species, but does not naturally infect the *Aedes aegypti* mosquitoes that transmit many important human arboviruses. When *Wolbachia* are artificially introduced, *Ae. aegypti* can be productively infected by the bacteria. Different *Wolbachia* strains can manipulate various aspects of mosquito biology in ways that are currently being exploited to mitigate the spread of *Aedes aegypti*-transmitted arboviruses (1). The two most important are cytoplasmic incompatibility (CI) and pathogen blocking (PB). PB is the restriction of viral infection and replication in mosquito tissues and thus the reduced likelihood of transmission of arboviruses by *Wolbachia*-infected mosquitoes. CI is a reproductive manipulation critical for the introgression of *Wolbachia* into natural mosquito populations because it reduces fertility when *Wolbachia*-infected males mate with uninfected females or with females infected with a different *Wolbachia* strain. Thus, it acts as a natural form of gene drive. For mitigation strategies that are dependent on PB, CI is harnessed to replace arbovirus-susceptible *Ae. aegypti* populations with a *Wolbachia-*infected *Ae. aegypti* population that has a reduced capacity to transmit arboviruses through PB—this is referred to as population replacement. The population replacement concept has advanced from laboratory experiments demonstrating the PB phenotype in *Ae. aegypti* for multiple arboviruses (2–7) to medium- to large-scale field deployments evaluating the efficacy of population replacement in at least 13 locations in Latin America, Asia, and the Pacific by the World Mosquito Program (WMP; https://worldmosquitoprogram.org) or *Wolbachia* Malaysia (https://imr.gov.my/wolbachia). Indeed, a randomized controlled trial conducted in Yogyakarta City, Indonesia, recently demonstrated that *Wolbachia* deployments reduced dengue incidence by 77% and dengue hospitalizations by 86% (8). In addition, it has been estimated that dengue incidence decreased by 96% at the original WMP release site in Cairns, Australia (9). WMP Brazil also estimated a 38% reduction in dengue cases and a 10% reduction in chikungunya virus infection incidence in Rio de Janeiro, where deployments began in 2017 (10). *Wolbachia* Malaysia estimated a dengue case reduction of 40.3% across all intervention sites (11).

Notably, the molecular mechanisms underlying the PB phenotype are poorly understood. However, the growing consensus in the field is that PB is complex, likely multifactorial, and likely context-dependent, meaning that the mechanisms controlling PB probably vary depending on the specific host–*Wolbachia* strain–virus combination (reviewed in (12)). As *Ae. aegypti* has no native *Wolbachia* symbionts, priming of the mosquito’s innate immune response has been previously implicated as a potential hypothesis to explain the PB phenotype of *w*Mel-infected *Ae. aegypti* (13–16), but the overall importance of innate immune priming remains unclear.

Non-canonical immune factors may also contribute to PB (17). For example, a recent study demonstrated that reduction in viral blocking in *w*Mel-infected *Ae. aegypti* was associated with reduced expression of a cell-cell adhesion gene (18). *Wolbachia* infection also promotes the induction of reactive oxygen species, which can directly kill pathogens through oxidative damage (15). Competition between arboviruses and *Wolbachia* for host resources has also been proposed as a mechanism for PB in insects, including mosquitoes (19,20). In sum, mosquitoes have complex responses to *Wolbachia* infection, which may involve many biological processes. As a result, PB could function through a mixed strategy that involves innate immune priming, oxidative stress, competition for key molecules like lipids, structural changes to the cellular environment, RNA translation, and/or cell adhesion (21,22), singly or in combination to limit virus replication.

*Ae. aegypti* harboring the *w*Mel strain of *Wolbachia* were first released in the field in Cairns, Australia in 2011. A recent study comparing *w*Mel genomes pre-release and nine years post-release demonstrated that the PB phenotype is remarkably stable and that releases of *Wolbachia*-infected mosquitoes for population replacement are likely effective for many years (23). However, it is important to note that *Wolbachia*-mediated suppression of virus infection is not complete (2,3,24), and therefore one might expect viruses to evolve resistance to this suppression. However, a few previous studies have suggested that it might be difficult for escape phenotypes to evolve (25–27), but without a better understanding of the selective pressures—that is the mechanisms underlying *Wolbachia* suppression—it is hard to analyze viral evolution or evaluate escape potential. Therefore, we sought to study both the mechanisms of *Wolbachia* suppression and the potential for virus evolution under *Wolbachia* selection.

Accordingly, we initiated transcriptome profiling studies to assess the whole set of molecular interactions involved in the dynamic processes of Zika virus (ZIKV) replication in *w*Mel-infected *Ae. aegypti* from the World Mosquito Program release site in Medellin, Colombia. We report that mosquito gene transcription dynamics exhibited temporal and tissue-specific expression patterns, and we contextualize this in relation to ZIKV infection dynamics in these same anatomical compartments. To investigate the impact of *Wolbachia* on ZIKV evolution, we used whole-genome deep sequencing to determine the intrahost genetic diversity of molecularly barcoded ZIKV populations replicating within *w*Mel-infected *Ae. aegypti* and *w*Mel-free *Ae. aegypti.* We show that ZIKV populations replicating in *Wolbachia*-infected mosquitoes or *Wolbachia*-free mosquitoes display signals of weak purifying selection and loose anatomical bottlenecks. The results from these evolutionary analyses suggest that weak, non-specific selection pressures inhibit the potential for *Wolbachia*-evading variants to emerge.

## Methods

### Ethics statement

This study was approved by the University of Minnesota, Twin Cities Institutional Animal Care and Use Committee (Protocol Number 1804–35828).

### Cells and viruses

African green monkey cells (Vero; ATCC CCL-81) and human embryonic kidney cells (HEK293T; ATCC CRL-3216) were cultured in Dulbecco’s modified Eagle medium (DMEM; Gibco) supplemented with 10% fetal bovine serum (FBS; Cytiva HyClone), 2 mM l-glutamine, 1.5 g/liter sodium bicarbonate, 100 U/ml penicillin, and 100 μg/ml streptomycin at 37°C in 5% CO2. The barcoded ZIKV infectious clone was constructed by bacteria-free cloning of the ZIKV PRVABC59 strain genome (GenBank accession no. KU501215.1), as previously described (28–30). The ZIKV PRVABC59 genome was amplified from an existing infectious clone (PMID: 27795432). The genetic barcode, with degenerate nucleotides at the third position of 8 consecutive codons in NS2A (30), was then introduced via an overlapping PCR-amplified oligonucleotide. The amplicons were assembled with a 5′ cytomegalovirus (CMV) promoter amplified from pcDNA3.1 (Invitrogen) by Gibson assembly (New England Biolabs), followed by enzymatic digestion of the remaining single-stranded DNA and noncircular double-stranded DNA. Full-length ZIKV constructs were amplified using rolling circle amplification (repli-g minikit; Qiagen) and genomic integrity was verified by restriction digestion and Sanger sequencing. Infectious barcoded ZIKV (ZIKV-BC) rescue was performed in HEK293T cells.

### Mosquito strains and colony maintenance

*Aedes aegypti* used in this study were maintained at the University of Minnesota, Twin Cities, as previously described (30,31). Two lines of mosquitoes were used in this study. The *Wolbachia-*infected *Aedes aegypti* line (COL.*w*Mel; infected with the wMel strain of *Wolbachia pipientis*) was established from several hundred eggs provided by the World Mosquito Program in Colombia (see https://www.worldmosquitoprogram.org/en/global-progress/colombia). Mosquitoes used in this study were from generations 7 to 14 of the laboratory colony. Wolbachia infection status was routinely verified using PCR with primers specific to the IS5 repeat element (4). The COL.*w*Mel line was cleared of *w*Mel infection at F3 by providing the colony with 1 mg/ml tetracycline solution (final concentration) dissolved in 10% sucrose *ad libitum*. Mosquitoes were treated with tetracycline as described in (32). Importantly, tetracycline-treated mosquitoes were allowed to recover for at least two generations, gut microflora was re-established, and mosquitoes were confirmed to be cured of *w*Mel as described above before being used in experiments. The tetracycline-treated line of mosquitoes is designated COL.tet.

### Vector competence and mosquito collection

Mosquitoes were exposed to ZIKV-BC by feeding on ketamine/xylazine-anesthetized ZIKV-BC-infected *Ifnar1^-/-^* mice, which develop sufficiently high ZIKV viremia to infect mosquitoes (2,33,34). *Ifnar1^-/-^* mice on the C57BL/6 background were bred in the pathogen-free animal facilities of the University of Minnesota, Twin Cities Veterinary Isolation Facility. 6 male and 3 female five- to seven-week-old mice were used for mosquito exposures. Mice were inoculated in the left hind footpad with 1 x 10^3^ PFU of ZIKV-BC in 25 µl of sterile PBS. Three- to six-day-old female mosquitoes were allowed to feed on mice two days post-inoculation. Subsequently, sub-mandibular blood draws were performed, and serum was collected to verify viremia. These mice yielded an infectious bloodmeal concentration of 4.2-4.8 log10 PFU per ml. Control mosquitoes were exposed to anesthetized, uninfected *Ifnar1^-/-^* mice. Mosquitoes that fed to repletion were randomized, separated into cartons in groups of 10–50, and maintained on 0.3 M sucrose in a Conviron A1000 environmental chamber at 26.5 ± 1 °C, 75% ± 5% relative humidity, with a 12 h photoperiod within the Veterinary Isolation Facility BSL3 Insectary at the University of Minnesota, Twin Cities. Infection, dissemination, and transmission rates were determined for individual mosquitoes, and sample sizes were chosen using long-established procedures (2,3,31). All samples were screened by plaque assay on Vero cells. Dissemination was indicated by virus-positive legs. Transmission was defined as the release of infectious virus with salivary secretions, i.e., the potential to infect another host, and was indicated by virus-positive salivary secretions. Midgut and carcass tissues were collected at 4 and 7 days post-blood feeding from additional mosquitoes exposed to the same infected or uninfected bloodmeals used for vector competence experiments. For each time point, 30 midguts and 30 carcasses were collected per strain per condition. Dissected tissues were immediately flash-frozen in microfuge tubes on dry ice and stored at −80°C until RNA isolation. Two complete sets of replicate samples (n=3 biological replicates total) were collected using distinct generations of mosquitoes to account for stochastic variations.

### Virus titration

Infectious virus was titrated by plaque assay on Vero cells. A confluent monolayer of Vero cells was inoculated with a 10-fold dilution series of each sample in duplicate. Inoculated cells were incubated for 1 h at 37°C and then overlaid with a 1:1 mixture of 1.2% Oxoid agar and 2× DMEM (Gibco) with 10% (vol/vol) FBS and 2% (vol/vol) penicillin-streptomycin. After 3 days, the cell monolayers were stained with 0.33% neutral red (Gibco). Cells were incubated overnight at 37°C, and plaques were counted. Plaque counts were averaged across the two replicates, and the concentration of infectious ZIKV-BC was back-calculated from the mean.

Viral RNA was isolated directly from mosquito bodies, legs, and saliva. Mosquito bodies and legs were homogenized in phosphate-buffered saline (PBS) supplemented with 20% FBS and 2% penicillin-streptomycin with 5-mm stainless steel beads with a TissueLyser (Qiagen) before RNA isolation. RNA was isolated with the Maxwell RSC viral total nucleic acid purification kit on a Maxwell RSC 48 instrument (Promega). Isolated ZIKV RNA was titrated by qRT-PCR using TaqMan Fast virus 1-step master mix (ThermoFisher) and a LightCycler 480 or LC96 instrument (Roche). Final reaction mixtures contained 600 nM each ZIKV-specific qRT-PCR primer (5′-CGY TGC CCA ACA CAA GG-3′ and 5′-CCA CYA AYG TTC TTT TGC ABA CAT-3′) and 100 nM probe (5′-6-carboxyfluorescein-AGC CTA CCT TGA YAA GCA RTC AGA CAC YCA A-black hole quencher 1-3′). Cycling conditions were 50°C for 5 min, 95°C for 20 s, and 50 cycles of 95°C for 15 s followed by 60°C for 1 min. ZIKV RNA titers were interpolated from a standard curve of diluted in vitro-transcribed ZIKV RNA. The limit of detection for this assay is 150 ZIKV genome copies/ml.

### Mosquito RNA Isolation

Total RNA was isolated from mosquito midguts and carcasses using the MasterPure RNA Purification Kit (Epicenter, Madison, WI), which included built-in Proteinase K and DNase treatments. Fisherbrand™ RNase-Free Disposable Pellet Pestles were used for tissue homogenization. RNA purity and concentration were determined using a Qubit 4 fluorometer (ThermoFisher) and further confirmed by Agilent 4200 TapeStation (Agilent Technologies, Santa Clara, CA). Only quality intact RNA was used for RNA sequencing.

### Illumina RNAseq library preparation and sequencing

Multiplex sequencing libraries were generated from 500 ng of total RNA (per library) using Illumina’s TruSeq sample prep kit and multiplexing sample preparation oligonucleotide kit (Illumina Inc., San Diego, CA) following the manufacturer’s instructions. Equal amounts of total RNA were pooled from replicates before library construction for midguts and carcasses to create 3 pools per strain per condition for each biological replicate. Samples were sequenced on an Illumina NovaSeq, which generated 2×150 bp paired-end reads at a depth of 20 million reads. Illumina’s bcl2fastq v2.20 was used for de-multiplexing, and sequence quality was assessed based on %GC content, average base quality, and sequence duplication levels.

### Sequence alignment and transcript quantification

RNA sequencing data were quality-checked using FastQC (v0.11.9) (35) and summarized using MultiQC (v1.12) (36). The resulting trimmed reads were aligned to the *Aedes aegypti* LVPm_AGWG reference genome (VectorBase: AaegL5.3) using kallisto (v0.46.1) (37), which relies on a pseudoalignment framework. Out of 3.7 billion sequence reads, 73-93% of reads mapped unambiguously to the *Ae. aegypti* reference genome (VectorBase: AaegL5.3). Downstream analysis followed the DIY Transcriptomics (38) R workflow, supplemented by bespoke R (v4.2.2) scripts (39). Aligned reads were annotated using the tximport (v1.28.0) R package (40). Differentially expressed genes were identified using raw gene counts, and heatmap analyses were performed using vst transformed gene counts. Differential gene expression analysis was performed using the DESeq2 (v1.40.1) package (41) using a significance cutoff of 0.01 and a fold change cutoff of log_2_ ≥ ±1. Volcano plots, temporal plots, and heatmaps were generated using the ggplot2 (v3.4.2) package (42) in R. Gene Set Enrichment Analysis was performed using GSEA (v4.3.2) (43,44) on normalized data against five lists of *Ae. aegypti* gene sets that were curated from previous genomic and transcriptomic studies (described in detail here (45)). Gene Ontology analysis was performed using the python (v3.11) package GO-Figure! (v1.0.1) (46). All data processing and analysis scripts are publicly available on GitHub (https://github.com/aliotalab/wMel-RNAseq-ms).

### Library preparation and Zika virus sequencing

Virus populations replicating in mosquito tissues were sequenced using a previously described tiled PCR amplicon approach (28,47,48). Briefly, a range of 800 - 6 x 10^6^ ZIKV genome copies/µL were converted to cDNA with the SuperScript^TM^ IV VILO^TM^ Master Mix that uses Superscript IV reverse transcriptase and random hexamer primers (ThermoFisher). PCR amplification of the entire ZIKV coding region was then performed in two reactions with pools of nonoverlapping PCR primer sets. PCR products were tagged with the Illumina TruSeq Nano HT kit. All libraries were quantified by a Qubit 3 fluorometer (ThermoFisher), and an Agilent Bioanalyzer assessed quality before sequencing with a 2 x 250 kit on an Illumina MiSeq. Additionally, a reference stock ZIKV of strain PRVABC59 was included in every sequencing run.

### Bioinformatic analysis of deep-sequencing data

For virus sequence data, a pipeline was generated to process raw Illumina reads, align reads at a normalized depth, call variants, and calculate diversity metrics as described in (28). Briefly, raw paired-end reads were adapter trimmed (cutadapt 3.4 (49) with Python 3.9.14 (50)), and then any paired-end reads with mismatched bases or less than a 50-bp overlap were filtered before error correction and merging (BBMap version 38.95 (51)). Next, merged reads were quality-trimmed to exclude 5’ and 3’ ends under Phred quality score 35, and primer sequences from the tiled primer sets were trimmed from the ends of the high-quality merged reads. Reads were normalized to a 31-mer depth of 2000 reads (samtools and htslib 1.9 (52); BBMap version 38.95 (51)) and aligned to the ZIKV PRVABC59 reference (bwa 0.7.17-r1188 (53)) with mismatch penalty 10. Variants with a Phred quality score ≥ 30 and coverage ≥ 300 reads were called against the PRVABC59 reference sequences with LoFreq* (version 2.1.5) (54) and annotated with SnpEff (55). Genome-wide and sliding-window nucleotide diversity (π, πN, and πS) were calculated with SNPGenie (v3; 2019.10.31) (56). SNPGenie analyses utilized variants of all frequencies, while all other iSNV analyses excluded variants at or below 1% allele frequency. For barcode analysis, high-quality merged reads were aligned to the ZIKV PRVABC59 reference sequence, and reads fully covering the barcode region were isolated. Barcode sequences were then extracted from the reads, and the abundance and species richness of unique barcodes was calculated as adapted from (28)). Barcode species richness was calculated as the number of unique barcode species occurring within an individual sample in RStudio (R version 4.2.1 “Funny-Looking Kid” and RStudio version “Spotted Wakerobin”). All data processing and analysis scripts are publicly available on GitHub (https://github.com/RiesHunter/Wolbachia).

### Statistical analysis

All statistical analyses from the transcriptomic data were conducted using GraphPad Prism 9 (GraphPad Software, CA, USA). Statistical analyses from the viral deep-sequencing data were conducted in RStudio, under the null hypothesis of equal means between groups with paired or unpaired Welch t-tests. Statistical significance was designated to P-values of less than 0.05.

### Data availability

Raw Illumina sequencing data are available on the NCBI Sequence Read Archive under BioProject no. PRJNA949154.

## Results and Discussion

### COL.wMel are resistant to Zika virus infection

To confirm that the phenotype of reduced vector competence existed in *w*Mel-infected *Ae. aegypti* on a Colombian genetic background (COL.*w*Mel) for our previously characterized barcoded ZIKV (ZIKV-BC; strain PRVABC59), we compared the relative abilities of COL.*w*Mel and COL.tet (tetracycline-treated mosquitoes to remove *Wolbachia*) to transmit ZIKV-BC in the laboratory. To assess vector competence, mosquitoes were exposed to viremic bloodmeals via feeding on ZIKV-BC-infected *Ifnar1^-/-^* mice. Infection, dissemination, and transmission rates were assessed at 4, 7, and 14 post-feeding (dpf) using an in-vitro transmission assay (2,31,33). ZIKV-BC infection status was confirmed by plaque assay to identify infectious virus. Consistent with our previous results using these mosquitoes (2), COL.*w*Mel displayed poor peroral vector competence for ZIKV-BC compared to COL.tet (**Fig 1**). Indeed, there was a significant reduction in ZIKV-BC infection status as compared to COL.tet at all time points and across all replicates (Fisher’s Exact test, 4 dpf: p < 0.0001, 7 dpf: p < 0.0001, 14 dpf: p < 0.0001), and ZIKV-BC was never detected in the saliva of COL.*w*Mel, thus confirming the PB phenotype in these mosquitoes.

**Fig 1.**
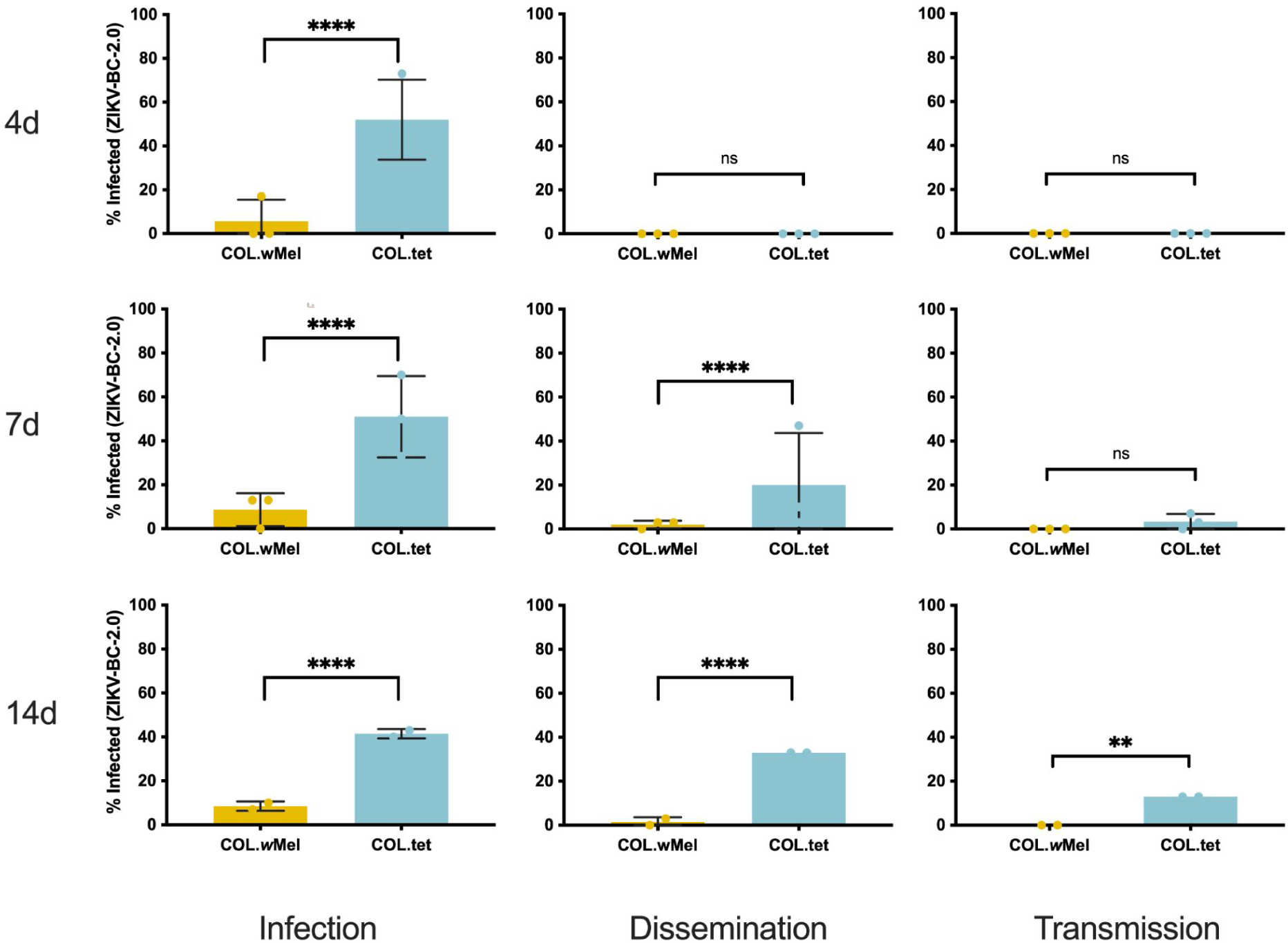
Vector competence of COL.*w*Mel and COL.tet orally exposed to ZIKV-BC. Mosquitoes were allowed to feed on ZIKV-BC-infected mice and were examined at days 4, 7, and 14 post-feeding to determine infection, dissemination, and transmission efficiencies. Infection efficiency corresponds to the proportion of mosquitoes with virus-infected bodies among the tested ones. Dissemination efficiency corresponds to the proportion of mosquitoes with virus-infected legs, and transmission efficiency corresponds to the proportion of mosquitoes with infectious saliva. *significant reduction in infection rates (Fisher’s Exact test *p < 0.05, **p < 0.01, ***p < 0.001, ****p<0.0001) **(A).** 4 days post-feeding (Biological replicate number 1, n = 30; Biological replicate number 2, n=30; Biological replicate 3, n=30) **(B).** 7 days post feeding (n = 30; Biological replicate number 2, n=30; Biological replicate 3, n=30) **(C).** 14 days post feeding (n = 30; Biological replicate number 2, n=No Data; Biological replicate 3, n=30).

### *Wolbachia* influences mosquito innate immune gene transcription

Next, we sought to determine the molecular basis of resistance to ZIKV-BC infection by comparing the transcriptional responses of COL.*w*Mel and COL.tet. We collected midgut and carcass tissue samples from COL.*w*Mel exposed to either a ZIKV-infected or uninfected bloodmeal. Midgut and carcass tissue samples were collected on days 4 and 7 dpf. These timepoints were chosen because they coincide with significant ZIKV infection in mosquitoes without *Wolbachia* (**Fig 1**). However, it is important to note that these timepoints may not have captured early responses that may also be important for PB. A matching set of tissue samples was concurrently collected from COL.tet to comparatively evaluate the transcriptome dynamics in a compatible mosquito-virus pairing. Poly(A)-selected mRNA isolated from carcasses and midguts were pooled and subjected to Illumina sequencing (142 libraries), and *Ae. aegypti* transcript quantification was carried out using kallisto (37). The overall ZIKV-BC infection prevalence ((PFU-positive + vRNA-positive)/total number of mosquitoes for all 3 biological replicates) was 81% for COL.*w*Mel and 93% for COL.tet on 4 dpf, respectively (**Fig S1**). First, because transcriptional responses are known to be altered in *Wolbachia*-infected mosquitoes compared to *Wolbachia*-free mosquitoes (57), we compared the transcriptional responses of COL.*w*Mel and COL.tet at 4 and 7 dpf that received an uninfected bloodmeal. There was a significant difference in the transcriptional responses of COL.*w*Mel versus COL.tet (Wald Test, p < 0.01) at 4 dpf, with 313 transcripts differentially regulated by twofold or greater in midguts and 927 in carcasses (**Fig 2A**). Of these, 75 (midguts) and 774 (carcasses) had an increased abundance and 238 (midguts) and 153 (carcasses) had a decreased abundance. Similarly, at 7 dpf, there were 596 transcripts differentially regulated by twofold or greater in midguts and 859 in carcasses. Of these, 127 (midguts) and 703 (carcasses) had an increased abundance and 442 (midguts) and 156 (carcasses) had a decreased abundance (**Fig 2A**). Since innate immune priming has been previously implicated as a potential hypothesis to explain the PB phenotype of *Wolbachia*-infected mosquitoes (13–16), we examined a set of 319 predicted immune genes (45) and found that 16% and 9.7% (51 and 31 transcripts) were differentially regulated in the carcasses at 4 and 7 dpf in COL.*w*Mel as compared to COL.tet; and 5.6% and 8.2% (18 and 26 transcripts) were differentially regulated in midguts at 4 and 7 dpf (**Fig 2B** and **Table S1**). Of these, the majority of carcass transcripts had increased abundance (95.1% or 82 transcripts total, 78 with increased abundance and 4 with decreased abundance), whereas the majority of midgut transcripts had decreased abundance (95.5% or 44 transcripts total, 2 with increased abundance and 42 with decreased abundance).

**Fig 2.**
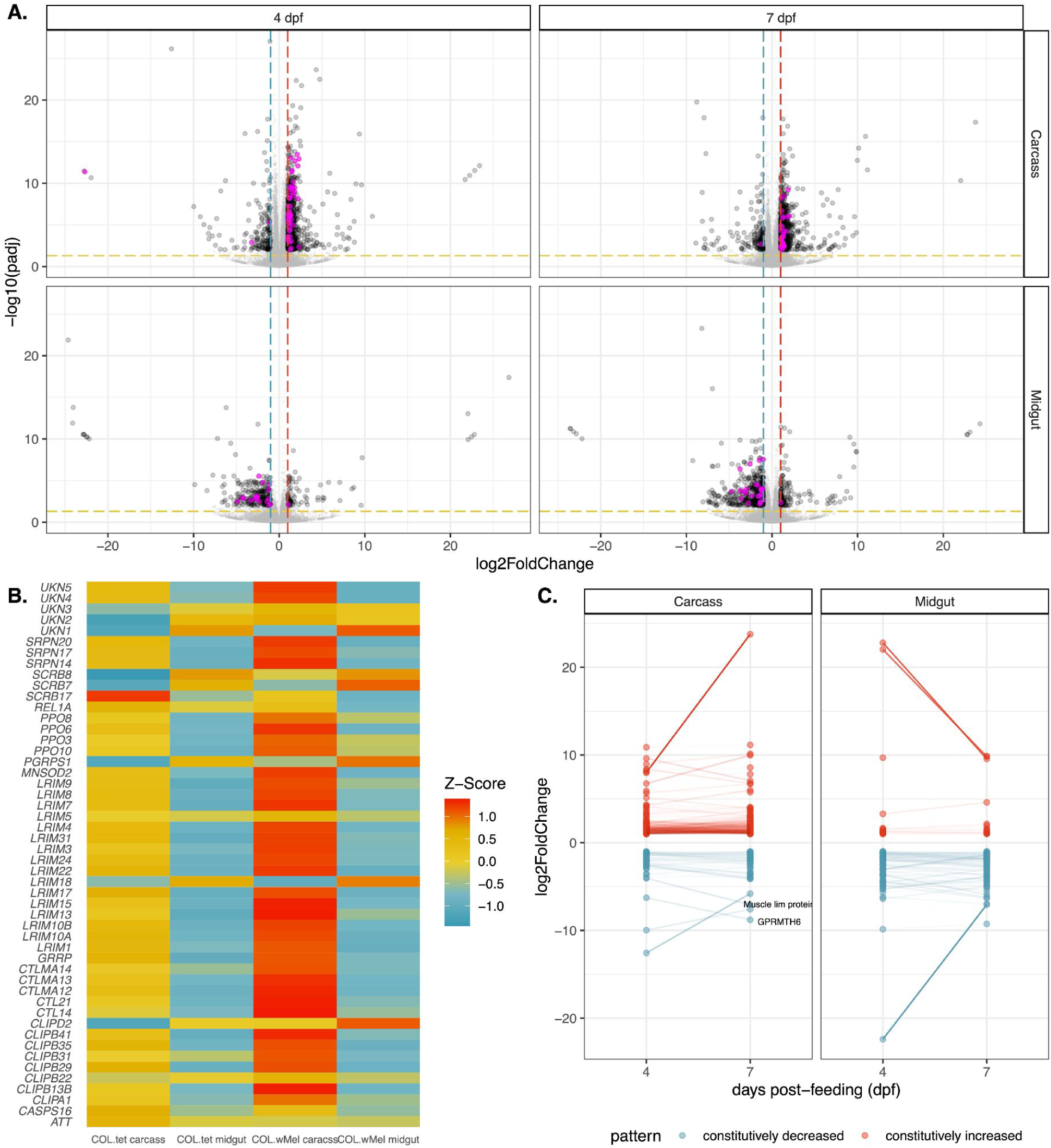
Transcriptional changes in COL.*w*Mel relative to COL.tet. **(A)** Volcano plots depicting differentially expressed transcripts in midgut and carcass tissues at 4 and 7 dpf, highlighting *Aedes aegypti* predicted innate immune genes (highlighted in magenta). Significant changes are transcripts with |log_2_FoldChange| >1 (vertical red and blue dashed lines) and -log10(padj) > 0.05 (horizontal yellow dashed line). **(B)** Heatmap of differentially expressed mosquito innate immune genes in COL.*w*Mel and COL.tet mosquitos that were exposed to an uninfected bloodmeal. The list of predicted innate immune genes was chosen from transcripts differentially regulated in COL.*w*Mel carcass tissue 4dpf. **(C)** Transcripts with constitutively increased abundance (red) and decreased abundance (blue) across both collection time points. Transcripts with a difference in log_2_FoldChange > 2.5 and that are annotated in VectorBase are labeled. Muscle lim protein (VB ID: AAEL019799). GPRMTH6 (VB ID: AAEL011521). No genes went from being significantly increased to significantly decreased or vice versa between 4 dpf and 7dpf.

To further characterize innate immune-related transcriptional activity between COL.*w*Mel and COL.tet, we used Gene Set Enrichment Analysis (GSEA) (43). Five gene sets were analyzed (45), including 1) predicted immune genes (45), 2) genes up-regulated by Toll, 3) by immune deficiency (Imd) (58), or 4) by Janus Kinase Signal Transducers and Activators of Transcription (JAK-STAT) signaling (59); and 5) genes that are up-regulated when mosquitoes are infected with *Wolbachia* (16). At 4 dpf, predicted immune genes and Imd gene sets were enriched in COL.*w*Mel relative to COL.tet in carcass tissue only (p-values = 0.032 and 0.028, respectively) (**Fig 2B** and **Table S1**). None of the gene sets were enriched at any other timepoint or in any other tissue type for this comparison. When we further examined the 51 immune transcripts differentially regulated in COL.*w*Mel relative to COL.tet carcass tissue at 4 dpf (**Fig 2B**), we found that there was 14% overlap between the predicted immune and Imd gene sets (REL1A, CLIPA1, CLIPB13B, CTLMA14, CTL14, UNK, and CTLMA12). Recent data suggest an antiviral role for nuclear factor kappa-light-chain-enhancer of activated B cells (NF-κB) signaling pathways— which are activated via the Imd pathway—against certain viruses in mosquitoes (reviewed in (60)). However, it has previously been shown that functional Toll and Imd pathways are not required for PB against DENV in *w*MelPop-infected *Drosophila* (61). Thus, PB is likely more complicated than a simple priming of the mosquito innate immune system, which is consistent with what others have reported previously (16,61).

### *Wolbachia* influences diverse cellular, biological, and molecular processes in the mosquito

To identify other factors that could be responsible for PB, we next performed Gene Ontology (GO) analysis to identify annotated functions that are overrepresented in the lists of transcripts that were differentially expressed in COL.*w*Mel compared to COL.tet. We focused our analysis on transcripts that were differentially expressed in the midgut at 4 dpf, because the midgut is the first anatomical barrier a virus encounters when infecting a mosquito, *Wolbachia* are present in this tissue (5); and we observed a significant reduction in overall ZIKV infection status at all timepoints, observed minimal evidence of disseminated infection, and did not detect virus in the saliva, all of which indicate reduced midgut susceptibility in COL.*w*Mel (**Fig 1**). The R package topGO was used to associate DEGs with GO terms, and to group semantically similar GO terms and reduce redundancy in topGO terms. We found that GO categories associated with muscle function, metabolism (both lipids and nucleotides), oxidative stress, and the cell cycle were overrepresented in the midgut at 4 dpf (**Fig 3**). These data suggest that the midgut environment of *Ae. aegypti* responds transcriptionally to the presence of *Wolbachia*, which is likely the result of the changing metabolic and energy needs required for *Wolbachia* endosymbiosis. We therefore postulate that PB results from immunological, physical, and/or physiological mechanisms directed at cellular damage, byproducts, or other factors resulting from infection with *Wolbachia*.

**Fig 3.**
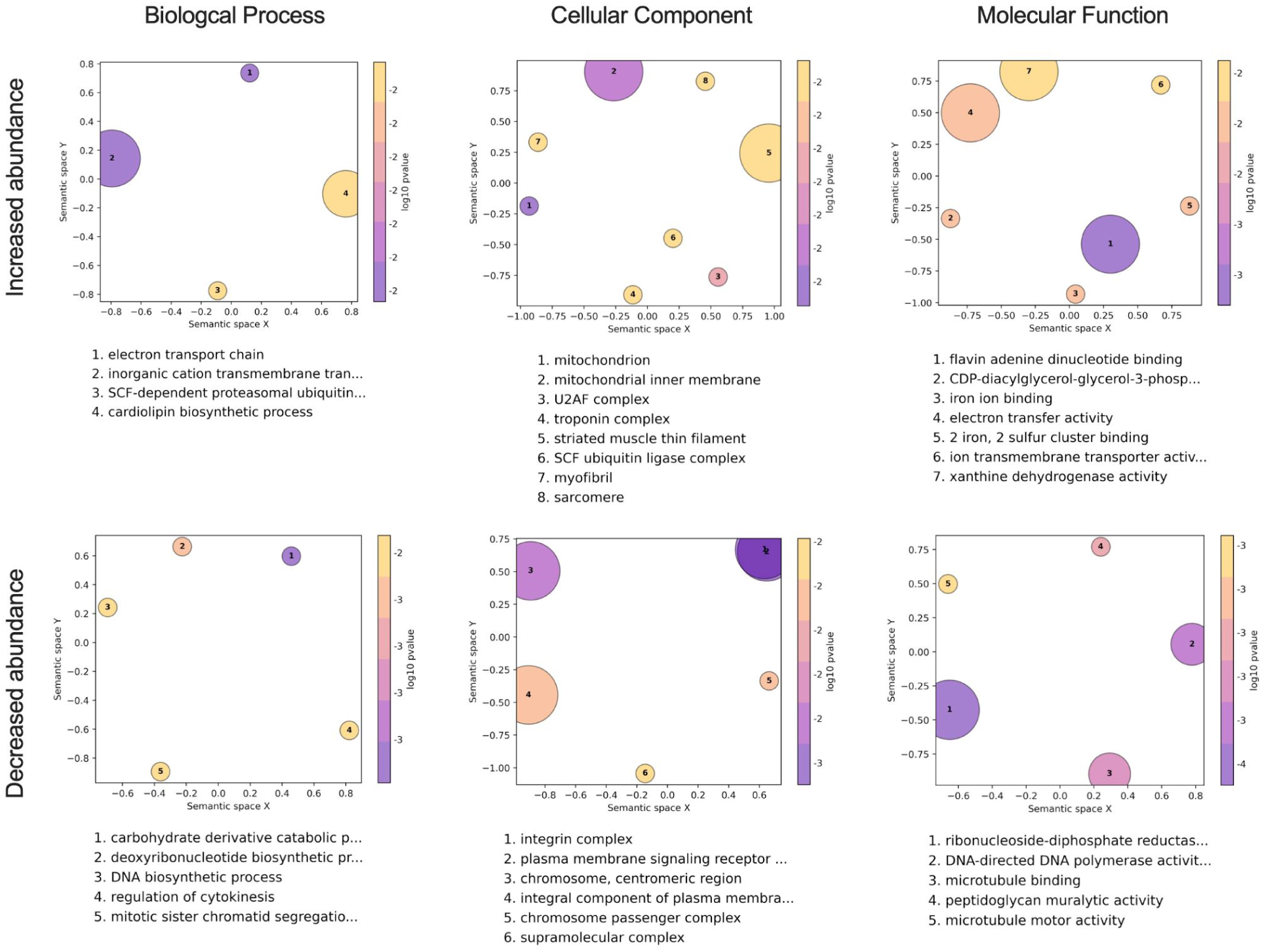
Gene Ontology analysis of COL.*w*Mel midguts 4 days post blood feeding. GO terms associated with the differentially expressed transcripts in COL.*w*Mel midguts at 4dpf. The top 10 GO terms from each category (Biological Process, Cellular Component, Molecular Function), determined by topGO, were run in the GO Figure! pipeline to combine semantically similar terms and reduce redundancy. Terms are ranked by lowest log10(p-value). The size of each graphical point corresponds to the number of topGO terms associated with the listed summarizing term.

For example, induction of transcripts involved in muscle function (GO categories: troponin complex, striated muscle thin filament, myofibril, and sarcomere) could be in response to the cellular damage and mechanical disruption the mosquito host experiences during *Wolbachia* infection (62–67). Similarly, Overrepresentation of the GO categories regulation of cytokinesis, mitotic sister chromatid segregation, chromosome, centromeric region, chromosome passenger complex, supramolecular complex, DNA-directed DNA polymerase activity, microtubule binding, microtubule motor activity, integrin complex, among others likely are the result of *Wolbachia’s* restructuring of the intracellular space (68) and could indicate that *Wolbachia* is making the cell a less hospitable environment for virus infection. Arboviruses use the cytoskeleton to enter/exit cells (69,70), so *Wolbachia*-induced modifications could be detrimental to viral replication.

Additionally, previous work has demonstrated that *Wolbachia* alters the host transcriptome at the interface of nucleotide metabolism pathways (71) and that there is competition between mosquito host and *Wolbachia* for lipids (20,72,73). Overrepresentation of GO categories involved in DNA biosynthetic process, deoxyribonucleotide biosynthetic process, carbohydrate derivative catabolic process, ribonucleoside-diphosphate reductase, cardiolipin biosynthetic process, integral component of plasma membrane, and CDP-diacylglycerol-glycerol-3-phosphate is consistent with these previous observations. As a result, PB could manifest because *Wolbachia* is dependent on the mutualistic relationship with its host to acquire all nutrients (66,67) and competition for resources could be detrimental to arbovirus infection.

Overrepresentation of the GO categories mitochondrion, mitochondrial inner membrane, iron ion binding, iron and sulfur cluster binding, the electron transport chain, electron transfer activity, and xanthine dehydrogenase are likely the result of the mosquito needing to manage oxidative stress, which also has been shown to coincide with *Wolbachia* infection (15). Mitochondria are also a source of reactive oxygen species (ROS), and ROS production is controlled by mitochondrial membrane potential (74). Production of ROS could result in PB through direct effects on the virus (75) or through downstream effects like activation of the Toll pathway to control virus infection (15). Mitochondrial function also controls epithelial barrier integrity and stem cell activity in the mosquito gut (76), again suggesting that the presence of *Wolbachia* may be creating a less hospitable environment for virus infection.

GO categories associated with metabolism, oxidative stress, cell signaling, and the cell cycle were also overrepresented in the midguts at 7 dpf, and in carcasses at 4 and 7 dpf (**Fig S2**). These data provide additional support for *Ae. aegypti* responding transcriptionally to the presence of *Wolbachia* in a temporal- and tissue-independent manner. That is, the changes required for the maintenance of *Wolbachia* endosymbiosis persist in multiple mosquito tissues without a distinct temporal pattern of induction among these genes (**Fig 2C**).

### ZIKV-BC infection does not induce a more robust innate immune response in COL.wMel compared to baseline

The previous comparisons establish that *Wolbachia* infection in COL.*w*Mel alters numerous physiological processes even in the absence of a pathogen, but the PB phenotype may require infection-induced transcriptional activity upon pathogen exposure to limit virus infection. To better compare the transcriptional responses of COL.*w*Mel and COL.tet, we examined gene expression dynamics in these mosquitoes exposed to ZIKV-BC. Four dpf, there was a significant difference in the transcriptional responses of COL.*w*Mel as compared to COL.tet (Wald Test, p < 0.01), with 81 transcripts differentially regulated by twofold or greater in midguts and 1,028 in carcasses (**Fig 4A**). Of these, 40 (midguts) and 871 (carcasses) had an increased abundance and 41 (midguts) and 157 (carcasses) had a decreased abundance. Similarly, at 7 dpf, there were 171 transcripts differentially regulated by twofold or greater in midguts and 692 in carcasses (**Fig 4A**). Of these, 79 (midguts) and 537 (carcasses) had an increased abundance and 92 (midguts) and 155 (carcasses) had a decreased abundance. Since immune response genes were found to be differentially regulated in mosquitoes with *Wolbachia* at baseline, we examined the set of 319 predicted immune genes (45,77) in the context of ZIKV-BC exposure, and found surprisingly few canonical immune genes differentially regulated in midguts at 4 (0.6% or 2 transcripts total; 1 with increased abundance and 1 with decreased abundance) or 7 dpf (0.3% or 1 transcript total; 1 with increased abundance and 0 with decreased abundance) in ZIKV-BC-exposed COL.*w*Mel relative to ZIKV-BC-exposed COL.tet. In carcasses, we found that 17.6% of immune genes (56 transcripts) were differentially regulated at 4 dpf and 8.2% (26 transcripts) at 7 dpf in ZIKV-BC-exposed COL.*w*Mel relative to ZIKV-BC-exposed COL.tet (**Fig 4A** and **Table S2**). We also performed GSEA using the same gene sets described earlier, but none of the gene sets were enriched at any timepoint or in either tissue type after ZIKV-BC infection. However, of the 56 immune genes that were differentially regulated in the carcass tissue at 4 dpf (**Fig 4B**), 72.5% overlapped with the list of 51 immune genes that were differentially regulated in carcasses at baseline (**Fig 2B**). These data suggest ZIKV infection-induced immune activation is not mediating PB in COL.*w*Mel.

**Fig 4:**
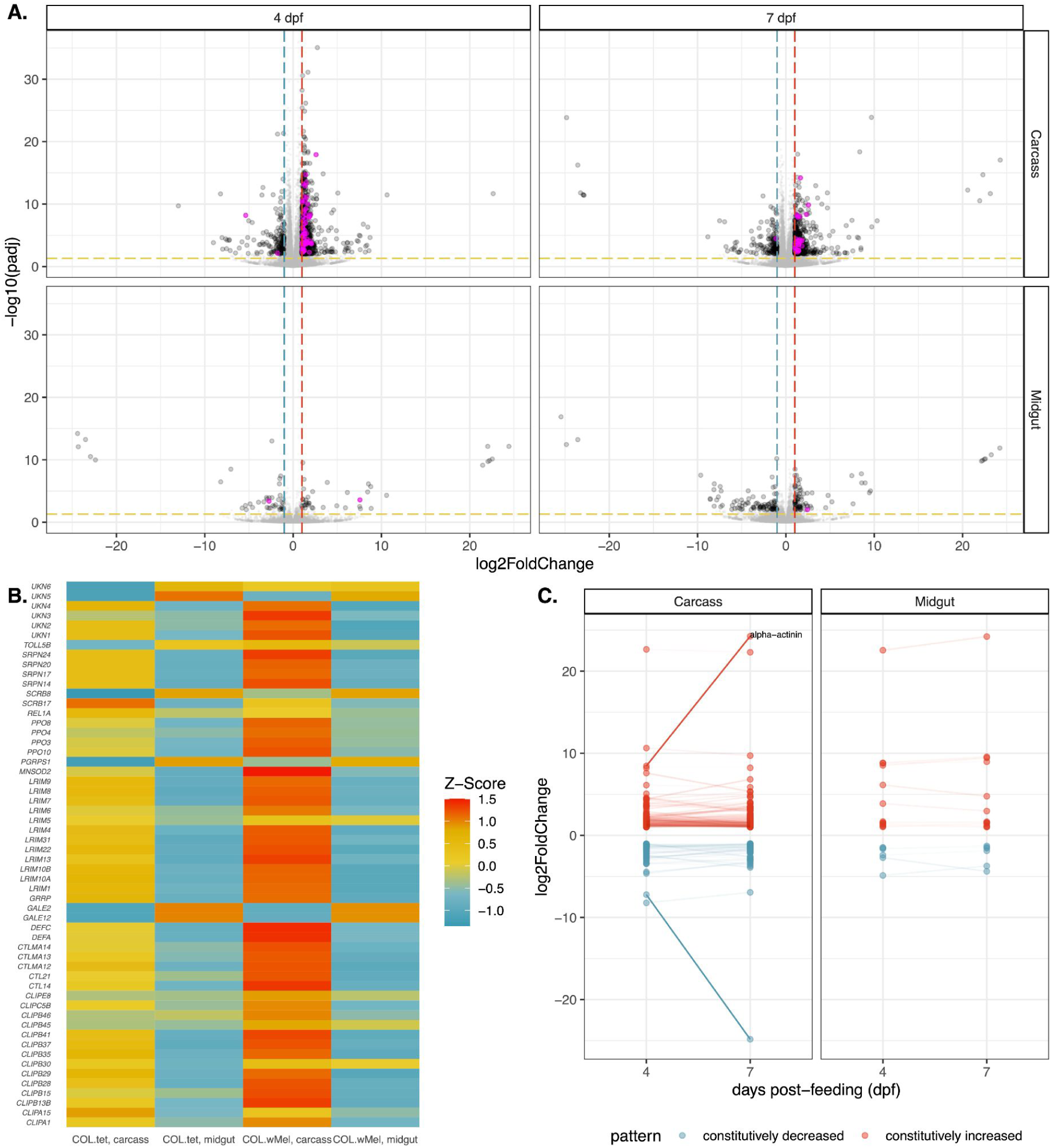
Transcriptional changes in COL.*w*Mel relative to COL.tet following a ZIKV-infected bloodmeal. **(A)** Volcano plots showing differentially expressed transcripts in midgut and carcass tissues at 4 and 7 dpf after a ZIKV-infected bloodmeal, highlighting *Aedes aegypti* predicted innate immune genes (highlighted in magenta). Significant changes are transcripts with |log2FoldChange| >1 (vertical red and blue dashed lines) and -log10(padj) > 0.05 (horizontal yellow dashed line). *Aedes aegpyti* predicted innate immune genes are highlighted in magenta. **(B)** Heatmap of mosquito innate immune gene expression patterns in ZIKV-exposed COL.*w*Mel and COL.tet mosquitos. The list of predicted innate immune genes was chosen from transcripts differentially regulated in COL.*w*Mel carcass tissue 4dpf. **(C)** Transcripts with constitutively increased abundance (red) and decreased abundance (blue) between both collection time points. Transcripts with a difference in log2FoldChange > 2.5 are labeled and that are characterized in VectorBase are labeled. Alpha-actinin (AAEL007306). No genes went from being significantly increased to significantly decreased or vice versa between 4 dpf and 7dpf.

As a result, we also performed GO analysis on COL.*w*Mel relative to COL.tet in the context of ZIKV-exposure, again focusing on the midgut tissue at 4 dpf. Similar to what we observed at baseline, we found GO categories associated with oxidative stress, metabolism, transcriptional pausing, cell membrane function/architecture, and signal transduction overrepresented in midguts at 4 dpf on a ZIKV-infected bloodmeal. These data suggest that the midgut environment of COL.*w*Mel responds transcriptionally in response to ZIKV exposure in a way that is similar to what occurs in response to *Wolbachia* alone. While it is possible that the factors involved in PB may not need to be induced upon pathogen exposure, PB likely involves multiple different biological processes in which *Wolbachia* infection alters the overall host environment of the mosquito in conjunction with infection-induced responses that also contribute to *Wolbachia*-mediated virus resistance. For example, the ubiquitin-proteasomal pathway has been shown to be important for flavivirus infection in mammalian and mosquito cells (78–81), and the overrepresentation of the GO terms SCF ubiquitin ligase complex, SCF-dependent proteosomal ubiquitination, positive regulation of apoptotic process, and positive regulation of cell death could indicate an antiviral state (**Fig 5**). Indeed, the ubiquitin/proteasome pathway plays an important role in controlling the level of programmed cell death (reviewed in (82)) and apoptosis has been known to have an antiviral role, including negatively impacting mosquito vector competence (65,83–88). SCF ubiquitin ligase also may be involved in triggering the innate immune response of *Ae. aegypti* and has previously been linked to the Toll pathway during Sindbis virus infection (89). Therefore, increased abundance of transcripts associated with the GO category SCF ubiquitin ligase complex could trigger increased Toll pathway activity to limit ZIKV infection.

**Fig 5.**
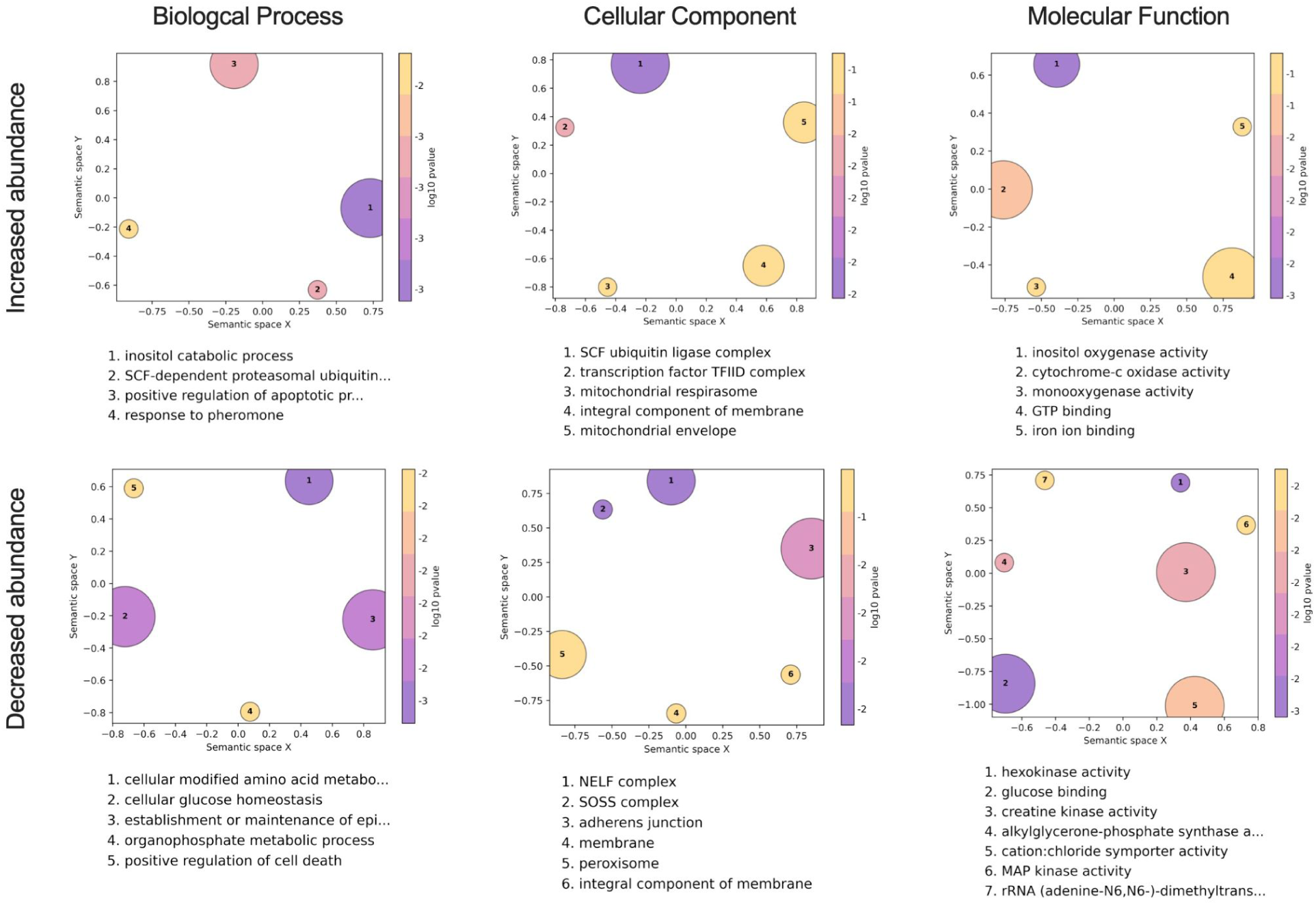
Gene ontology analysis in COL.*w*Mel midguts 4 days post feeding on ZIKV-infected mice. GO terms associated with the differentially expressed transcripts in COL.*w*Mel relative to COL.tet midguts at 4dpf on ZIKV-infected mice. The top 10 GO terms from each category (Biological Process, Cellular Component, Molecular Function), determined by topGO, were run in the GO Figure! pipeline to combine semantically similar terms and reduce redundancy. Terms are ranked by lowest log10(p-value). The size of each graphical point corresponds to the number of topGO terms associated with the listed summarizing term.

The role of transcriptional pausing in mosquito antiviral immunity remains unknown, but antiviral immunity in *Drosophila* requires transcriptional pausing to rapidly transcribe components of the RNA silencing, autophagy, JAK/STAT, Toll, and Imd pathways and multiple Toll-like receptors upon arbovirus infection (90). We found GO categories NELF complex, transcription factor TFIID complex, SOSS complex, and rRNA (adenine-N6, N6)-dimethyltransferase overrepresented in our dataset (**Fig 5**). These data could indicate that the COL.*w*Mel transcriptional pausing pathway is functioning to facilitate rapid induction of virus-induced genes and thus a robust antiviral response (91). Another notable feature of ZIKV-infected COL.*w*Mel midgut transcriptome was the overrepresentation of GO categories associated with oxidative stress, metabolism, and cell replenishment. Overrepresentation of GO categories associated with the same general biological, molecular, or cellular functions were observed in midguts and carcass tissue at both 4 and 7 dpf (**Figs 5 and S3**), similar to what we observed with COL.*w*Mel in the absence of ZIKV (**Figs 3 and S2**). Overall, these data suggest considerable change in the COL.*w*Mel transcriptome in response to ZIKV infection.

For each mosquito type (COL.*w*Mel and COL.tet), we also measured differential expression between ZIKV-infected relative to uninfected mosquitoes. We found that ZIKV infection induces dissimilar responses between mosquito types (**Fig 6**). The only timepoint and tissue type that had shared transcripts (n = 22) was the midgut tissues at 7 dpf (Pearson’s R^2^ = 0.104), and 45% of these were transcripts with unspecified annotations (10/22). These data suggest that ZIKV infection induces a distinct response in COL.wMel compared to COL.tet, and that response was dominated by the midgut tissue experiencing the most extensive transcriptional changes, with a large increase in differentially regulated transcripts from 4 dpf to 7 dpf: 26 to 371 transcripts respectively (**Fig 6**). The carcass tissue underwent modest transcriptional changes, with a similar increase in differentially regulated transcripts from 4 dpf to 7 dpf: 6 to 94 transcripts, respectively (**Fig 6**). Of the 371 and 94 transcripts that showed significantly different transcriptional activity at 7 dpf in the carcass and midgut tissues, respectively, 161 (43%) and 44 (47%) of those transcripts were unknowns, meaning that they have no previously described function. This suggests that many factors that could be involved in PB are not known (see **Tables S3** and **S4** for the full list of transcripts in these tissue types and at this timepoint).

**Fig. 6.**
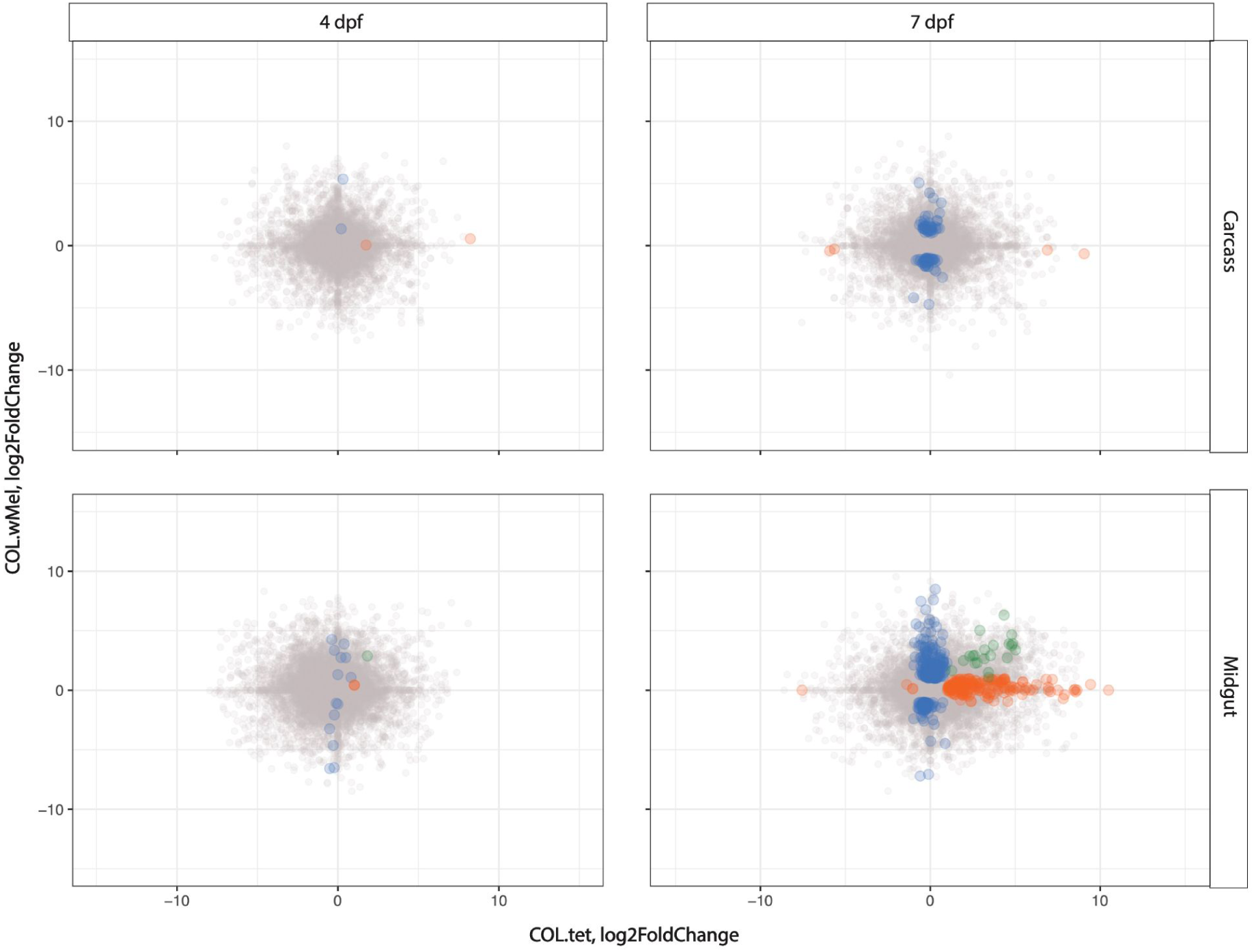
The induced response to ZIKV is distinct in COL.*w*Mel compared to COL.tet. The correlation in log_2_ fold change between COL.*w*Mel and COL.tet is shown at 4 and 7 days post feeding in carcass and midgut tissues. The transcripts highlighted in color differ in the direction or magnitude of expression (interaction term significant at p<0.01). Blue indicates transcripts unique to COL.*w*Mel, orange indicates transcripts unique to COL.tet, and green indicates transcripts that are shared between the two mosquito types.

### Gene transcription dynamics from ZIKV-BC infection in COL.tet

Next, we examined genome-wide temporal changes in transcript expression patterns in an effort to better understand the dynamic progression of ZIKV infection processes in the susceptible mosquito line, COL.tet. The midgut tissue underwent extensive transcriptional changes, with a large increase in differentially regulated transcripts from 4 dpf to 7 dpf: 12 to 378 transcripts, respectively (**Fig 7A**). In contrast, the carcass tissue underwent few transcriptional changes, with 6 transcripts differentially regulated at 4 dpf and 8 transcripts at 7 dpf. These data are somewhat surprising, since by 7 dpf, ZIKV-BC had disseminated from the initial site of replication in the midgut to secondary tissues—including salivary glands—throughout the mosquito body cavity in at least a subset of mosquitoes (**Fig 1**). Still, infection-induced gene expression changes exhibited temporal kinetics that appeared to closely reflect infection data reported in Fig 1. As a result, the mosquito’s transcriptional responses at these two timepoints likely reflect the long-term persistent effects of ZIKV-BC infection, some of which may represent a compensatory host response to cytopathic effects and other pathologic changes caused by intracellular virus replication (92,93) (**Fig 7B**). In addition, many cell-mediated defense responses in mosquitoes resemble basic cellular processes used in daily physiological processes and this may explain the increased abundance of transcripts associated with the GO categories reproduction, reproductive process, binding, mRNA binding, and protein binding (**Fig 7B**). Interestingly, there were only 8 transcripts (AAEL004386 chorion peroxidase; AAEL004388 heme peroxidase; AAEL004390 heme peroxidase; AAEL004978 DEAD box ATP-dependent RNA helicase; AAEL013063 autophagy related gene; AAEL013227 PIWI; AAEL014141 Serine Protease Inhibitor; and AAEL014251 Inhibitor of Apoptosis (IAP) containing Baculoviral IAP Repeat(s)) that overlapped with the set of 319 predicted immune genes (45). In addition, GSEA indicated that the JAK-STAT gene set was negatively correlated in midgut at 4 dpf and in the carcass tissue at 7 dpf (p = 0.047 and p = 0.039, respectively), and that the Toll gene set also was negatively correlated in carcasses at 7 dpf (p = 0.033). These data suggest that the transcriptional changes we observed may reflect the virus’s ability to persistently evade the host’s defenses to remain inside the host to achieve eventual transmission (94).

**Fig 7.**
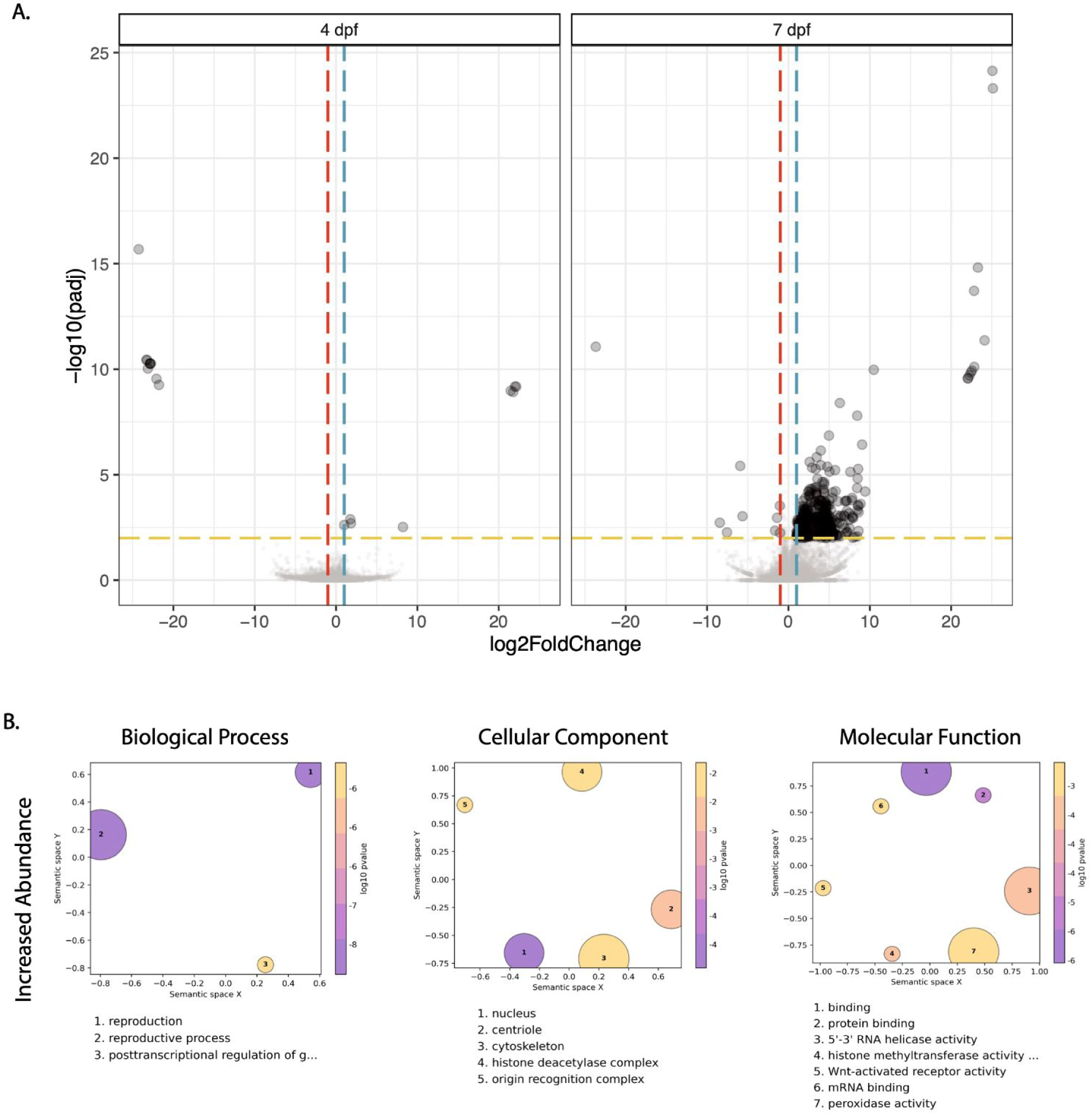
Differentially expressed transcripts and gene ontology analysis in COL.tet midguts 7 days post-feeding on ZIKV-infected mice. (A). Volcano plots demonstrating differentially expressed transcripts in the midgut of COL.tet midguts. Significant changes are transcripts with |log2FoldChange| >1 (vertical red and blue dashed lines) and -log10(p-adjusted) > 0.05 (horizontal yellow dashed line). **(B).** GO terms associated with transcripts with an increased abundance in COL.tet midguts 7dpf. Terms are ranked by lowest log10(p-value). The size of each graphical point corresponds to the number of topGO terms associated with the listed summarizing term.

### Zika virus populations are subject to weak purifying selection in both COL.*w*Mel and COL.tet mosquitoes

Blockage of ZIKV replication in COL.*w*Mel is incomplete, creating a scenario in which natural selection might favor the emergence of viral variants that can overcome PB. To address this possibility, we characterized the evolutionary dynamics of ZIKV-BC in the presence versus the absence of *Wolbachia*. First, we examined within-host viral genetic diversity, reasoning that it would be reduced in COL.*w*Mel compared to COL.tet mosquitoes. We identified intrahost single-nucleotide variants (iSNVs) present throughout the viral genome in the bodies (n=17 COL.*w*Mel and n=35 COL.tet), legs (n=2 COL.*w*Mel and n=19 COL.tet), and saliva (n=10 COL.tet) of mosquitoes that had fed on ZIKV-BC-infected mice. For comparison, we also sequenced viral genomes from two mice that were used to infect the mosquitoes. We could not recover additional blood samples from mice due to the large volume taken during mosquito feeding. Contrary to our expectations, viral nucleotide diversity (π) tended to be roughly equivalent in COL.*w*Mel compared to COL.tet across sampled time points (**Fig 8A**). For example, the number of iSNVs and the divergence among samples increased relative to the infected mice, which served as the source of mosquito viral populations, but were similar across all groups (**Fig S4**). We then estimated the strength and direction of natural selection operating on ZIKV within hosts by comparing levels of synonymous and nonsynonymous nucleotide diversity, denoted respectively as πS and πN. The levels of πN remained stable across anatomical sites and over time, while πS trended slightly downward over time in COL.tet bodies (**Fig 8A**). In general, πN - πS < 0 indicates that purifying selection is acting to remove deleterious mutations, while πN - πS > 0 signals that positive or diversifying selection is acting to favor new viral variants. Mean πN - πS values were consistently less than zero for all tissue types in COL.*w*Mel and COL.tet (**Fig 8B**), suggesting that virus populations were primarily under purifying selection. πN-πS differences were significantly lower from those observed in the infected mice across all time points, other than COL.*w*Mel legs, which did not have a normal data distribution due to low sample number (paired Welch t-test; p < 0.001). πN - πS was signficantly decreased between COL.tet body and COL.*w*Mel body, but comparisons between COL.tet legs and COL.*w*Mel legs could not be made due to low sample number (unpaired Welch t-test; p < .001). Additionally, the frequency-distribution spectrum of iSNVs was similar among all groups and supported the conclusion that ZIKV genomes were under weak purifying selection in all mosquitoes (**Fig S5**). We also calculated πN - πS differences by each gene in the ZIKV ORF, to determine whether natural selection might act differently on different genes.

**Fig 8.**
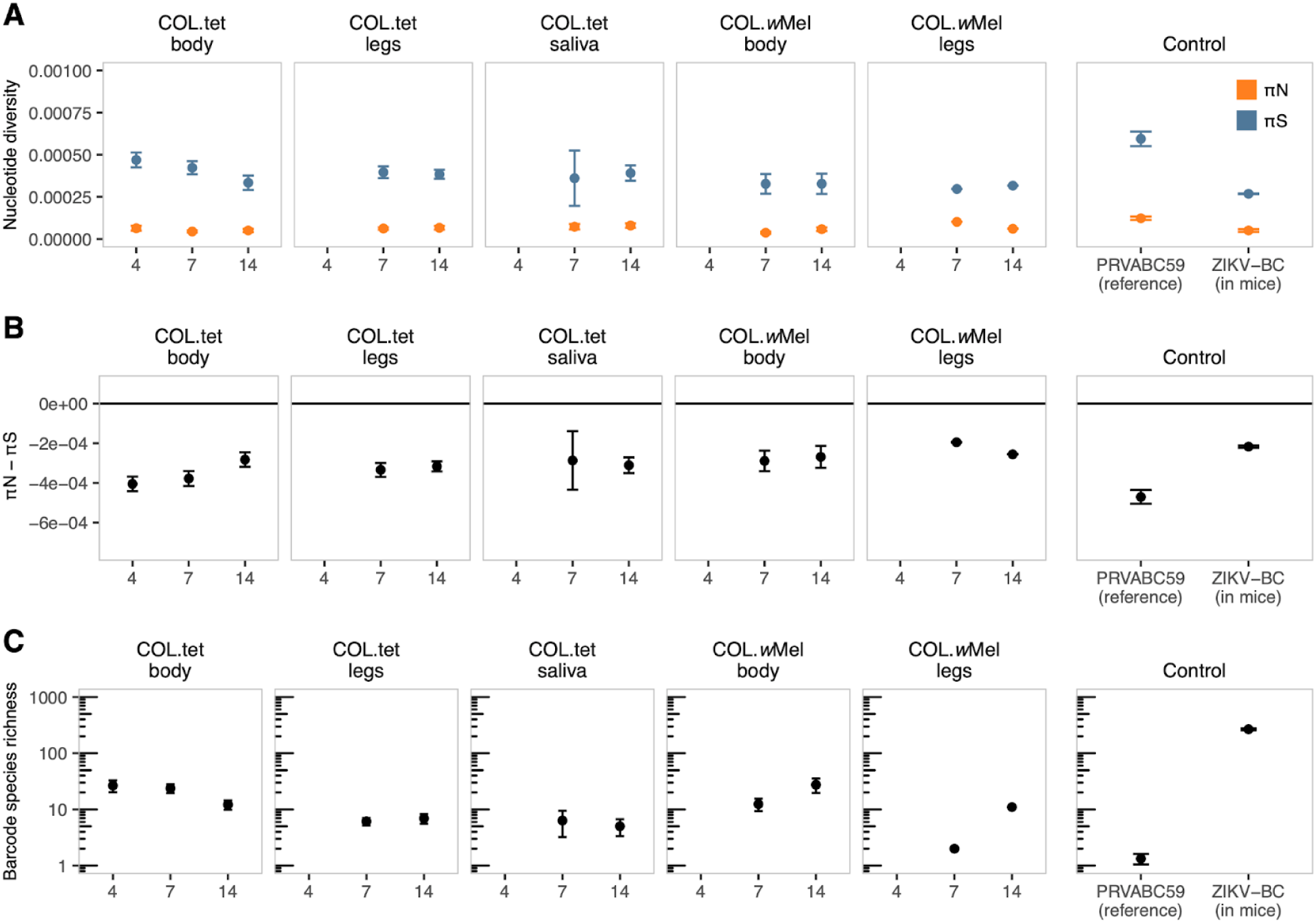
Zika virus is under weak purifying selection in *Wolbachia*-infected and *Wolbachia*-free mosquitoes. **(A).** Per-sample nucleotide diversity is quantified for nonsynonymous (πN; orange) and synonymous (πS; blue) sites across all ZIKV plaque-positive tissues collected from COL.*w*Mel and COL.tet mosquitoes. **(B).** The difference between πN and πS is plotted against the null hypothesis of neutral selection, denoted by a horizontal black line at zero. **(C).** Barcode species richness was quantified as the number of unique barcode species detected in each sample. All groups underwent 10,000 Bayesian bootstrap replicates, from which mean values and standard deviations were calculated.

For example, a trend of purifying selection across the genome could mask signs of positive selection focused on, say, the envelope gene. We found that results differed by gene, with some genes undergoing weak positive selection or mild purifying selection, while others showed no significant difference between the rates of nonsynonymous and synonymous substitutions (**Fig S6**). Together, these data suggest that selection pressures on the gene and genome level are weak and tend toward purifying selection. This is consistent with expectations for a relatively fit RNA virus replicating in its host and reflects a situation in which most mutations away from the consensus are neutral or mildly deleterious.

We next used the molecular barcode present in ZIKV-BC to assess the influence of anatomical bottlenecks within the mosquitoes in the presence and absence of *Wolbachia.* ZIKV-BC contains a run of eight consecutive degenerate codons in NS2A (amino acids 144 to 151), which generates significant viral diversity within the barcode region (28), and thus allows us to track the number and abundance of specific barcodes across anatomical compartments using deep sequencing. Consistent with our previous findings (28–30,34), the number of unique barcodes, which we will term barcode species richness, significantly declined over time in each anatomical compartment as the virus disseminated throughout the mosquito body cavity (unpaired Welch t-test; p < 0.001), with the lowest number of barcodes detected in mosquito saliva from COL.tet (**Fig 8C**). Statistical comparisons of barcode species richness from the COL.*w*Mel legs group could not be calculated due to small sample numbers. We also were not able to recover ZIKV from mosquito saliva from COL.wMel mosquitoes because effective PB prevented ZIKV dissemination to saliva. We detected an average of 267 unique barcodes in ZIKV-BC-infected mice compared to ∼10 unique barcodes in mosquito tissues, a significant reduction in barcode species richness (paired Welch t-test; p < 0.001). Barcode species richness could not be assessed in the COL.wMel legs group due to low sample number. We observed significantly higher barcode species richness in COL.tet bodies at 7 dpf relative to COL.*w*Mel bodies, while at 14 dpf we found the opposite: barcode species richness was higher in COL.wMel bodies than in COL.tet bodies (unpaired Welch t-test; p < 0.001). It is not possible that barcode species richness increased over time in mosquitoes, so the apparent increases observed in COL.wMel bodies and legs between days 7 and 14 are likely attributable to sampling error due to very small sample sizes. Taken together, these results suggest that barcode diversity is greatly reduced during mouse-to-mosquito transmission, consistent with prior work (28,30). Notably, we do not observe clear indications of barcode species richness decreasing through sequential within-host anatomical bottlenecks in mosquitoes, as some others have reported previously (29,95,96). The relative preservation of barcode species richness in mosquitoes in our study could reflect a greater ability to resolve within-host population sizes due to the artificially high sequence diversity conferred by the barcodes, but differences in experimental systems, sequencing, and analysis approaches may also contribute.

To help resolve these possibilities, we also investigated the fate of a synonymous iSNV in NS3 at genome position 5155 (**Fig S4**). As shown in the frequency-distribution spectra in **Fig S5**, iSNV C5155T is present in the stock sample of the reference virus (PRVABC59) and the ZIKV-BC samples at frequencies typically ranging from 30–50%. iSNV 5155T was responsible for much of the synonymous nucleotide diversity in whole-genome analyses. If within-host bottlenecks usually involve passage of 1-2 virions, we would expect this iSNV to be either stochastically fixed or lost in most samples. However, 5155T increased to >95% frequency in only one sample (COL.*w*Mel body at 14 dpf), while it declined to < 5% frequency in only 4 others (COL.tet saliva at 7 and 14 dpf; COL.tet body at 14 dpf; **Fig S7**). In all other samples (n=78), 5155T had a mean frequency of 40.8% and a median frequency of 38.3%, with a range of 20.4% to 74.7%. The preservation of both nucleotide variants at position 5155 in most samples therefore agrees with our finding that barcode richness is maintained through anatomical compartments in mosquitoes in our study and further suggests that in this system within-host bottlenecks typically involve passage of more than 1-2 virions. Consistent with this, a previous study showed that the complexity of West Nile virus populations replicating in mosquitoes was not significantly diminished by anatomical barriers (97).

### No evidence for positive selection for individual mutations within hosts

Our nucleotide diversity analyses suggest that *w*Mel does not exert strong selection pressures on ZIKV evolution. However, it is important to consider that natural selection may not act efficiently if effective population sizes are extremely small, which our data suggest, and gene- or genome-level summary statistics could miss selection for individual amino acid substitutions. To investigate this, we examined iSNVs that reached ≥ 50% allele frequency, i.e., the consensus level. We identified seventeen consensus-changing iSNVs, of which five were nonsynonymous, encoding amino acid changes NS1:N109S (57%), NS1:D180N (62%), NS2A:A117T (84%), NS5:L76R (56%), and NS5:I570L (99%). NS5:I570L and NS2A:A117T occurred in COL.*w*Mel bodies at 7 and 14 dpf, respectively. NS1:N109S and NS1:D180N were detected in COL.tet legs at 14 dpf, and NS5:L76R in COL.tet saliva at 14 dpf. None of these substitutions were observed in the ZIKV-infected mice or the stock reference virus controls, suggesting that they arose *de novo* in our samples. Small effective population sizes within infected mosquitoes could allow some mutations to rapidly change frequency, even if average within-host bottleneck sizes are greater than 1-2. Therefore we cannot conclude that these mutations reached high frequencies through the action of natural selection. In this regard it is notable that among the small number of consensus-changing within-host mutations, there was an excess of synonymous substitutions (12 of 17 consensus-changing mutations). Combining all our data, we find no evidence for positive selection for escape from *Wolbachia*-mediated PB during ZIKV replication in mosquitoes.

## Conclusions

Our findings suggest that the underlying mechanisms controlling *Wolbachia*-mediated PB likely involve many different immunological and physiological processes that function as part of a larger (and possibly redundant) network capable of limiting virus replication in the mosquito. It is also possible that PB may not be entirely regulated at the transcriptional level. Rather, the factors responsible could exist in the hemolymph or in another anatomical compartment in primed states—meaning that they do not return to basal levels after activation—as pre-PB complexes, thereby limiting or not requiring PB-related transcriptional activity upon virus exposure. Regardless, additional work will be required to disentangle the factors responsible. We also find that ZIKV is subject to weak purifying selection in mosquitoes in both the presence and absence of *Wolbachia*, similar to findings for RNA viruses in other hosts during acute infections. Using a barcoded ZIKV that allows us to sensitively quantify the number of distinct viral variants within mosquitoes, we find that within-host viral diversity is not significantly diminished in the presence vs. absence of *Wolbachia*, nor over the course of dissemination through mosquito tissues. Together our data suggest that PB is a complex and multifactorial phenotype, while natural selection is unlikely to act efficiently on ZIKV replicating in mosquitoes due to small within-host population sizes. These observations could help explain why we see little evidence for the emergence of *Wolbachia*-escape mutations. We posit that PB could be viewed as similar to combination antiretroviral therapy for HIV, in that using multiple complementary mechanisms to inhibit virus replication constrains the potential for resistance to evolve. Nonetheless our findings do not preclude the possibility that *Wolbachia* resistance could emerge over time, as arboviruses are passaged between humans and *Wolbachia*-infected mosquitoes in nature. Our study underscores the importance of both better defining the mechanisms of *Wolbachia*-mediated pathogen blocking and assessing the potential of viruses to evolve PB resistance to forecast the long-term sustainability of *Wolbachia* biocontrol.

## Acknowledgments

We acknowledge the University of Minnesota, Twin Cities BSL3 Program for facilities and Neal Heuss for support. We thank Elise Pritchard for her contribution to mosquito colony maintenance. We thank the University of Minnesota’s Genomics Center for RNA sequencing and the Minnesota Supercomputing Institute for computing resources. We also thank the World Mosquito Program for sharing *Wolbachia*-infected mosquitoes.

Research reported in this publication was supported by the National Institute of Allergy and Infectious Diseases of the National Institutes of Health under award number R21AI131454 and R01 AI132563 to M.T.A. A.S.J. is supported by the University of Minnesota, Twin Cities, Institute for Molecular Virology Training Program predoctoral fellowship number T32AI083196. Virus sequencing performed at the Wisconsin National Primate Research Center is supported by grants P51RR000167 and P51OD011106. The publication’s contents are solely the responsibility of the authors and do not necessarily represent the official views of the NIH.

## Notes

https://github.com/aliotalab/wMel-RNAseq-ms

https://github.com/RiesHunter/Wolbachia

### Competing Interest Statement

The authors have declared no competing interest.

