## Supplemental Figures for "*Wolbachia*-mediated resistance to Zika virus infection in *Aedes aegypti* is dominated by diverse transcriptional regulation and weak evolutionary pressures"

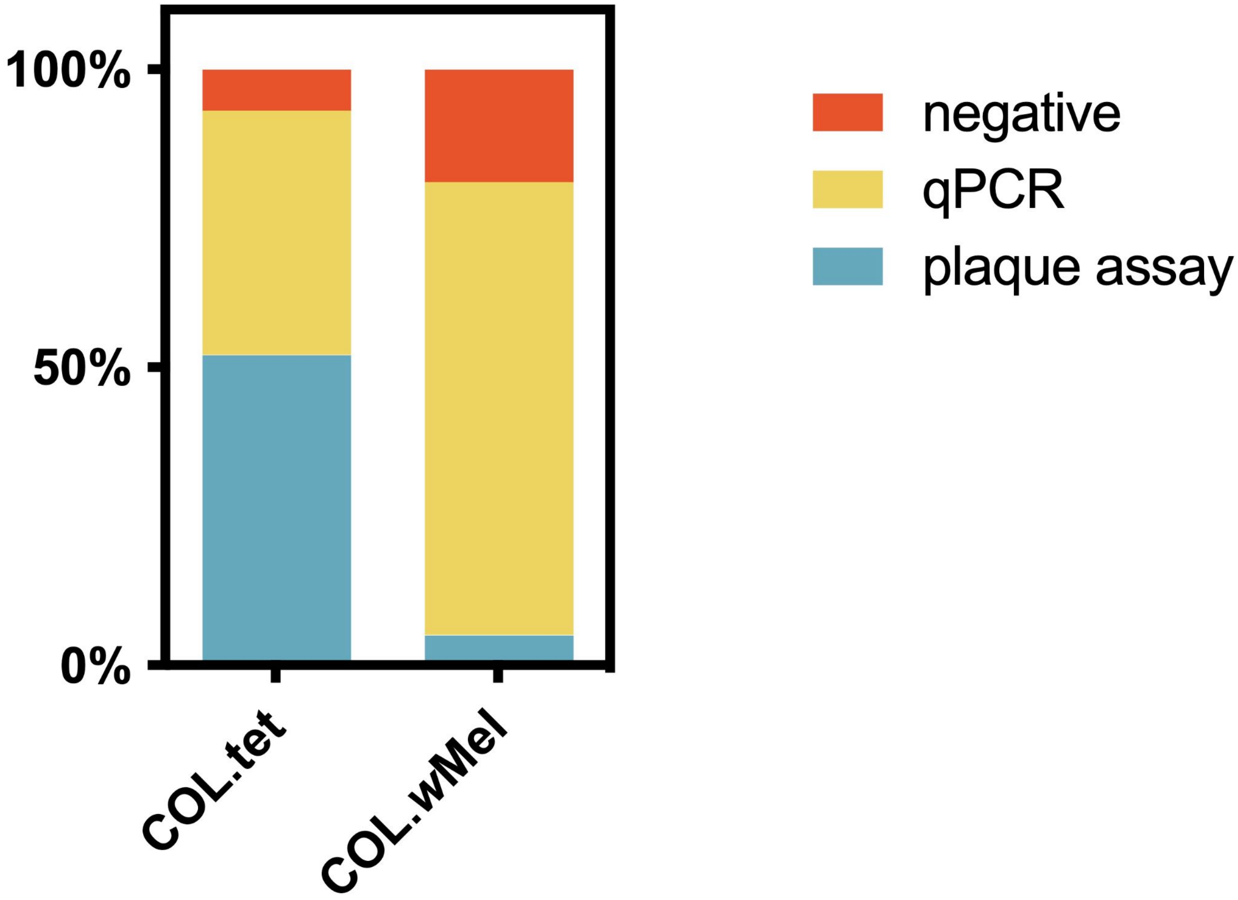


**Supplemental Figure 1: ZIKV infection prevalence in mosquito samples collected at 4 dpf.** Infection prevalence was measured by combining the number of PFU-positive mosquito samples via plaque assay (blue) and vRNA-positive samples via qPCR (yellow). Overall infection prevalence was 81% for COL.*w*Mel and 93% for COL.tet. Overall infection prevalence was calculated by adding PFU-positive and vRNA positive samples and dividing by the total number of mosquitoes.

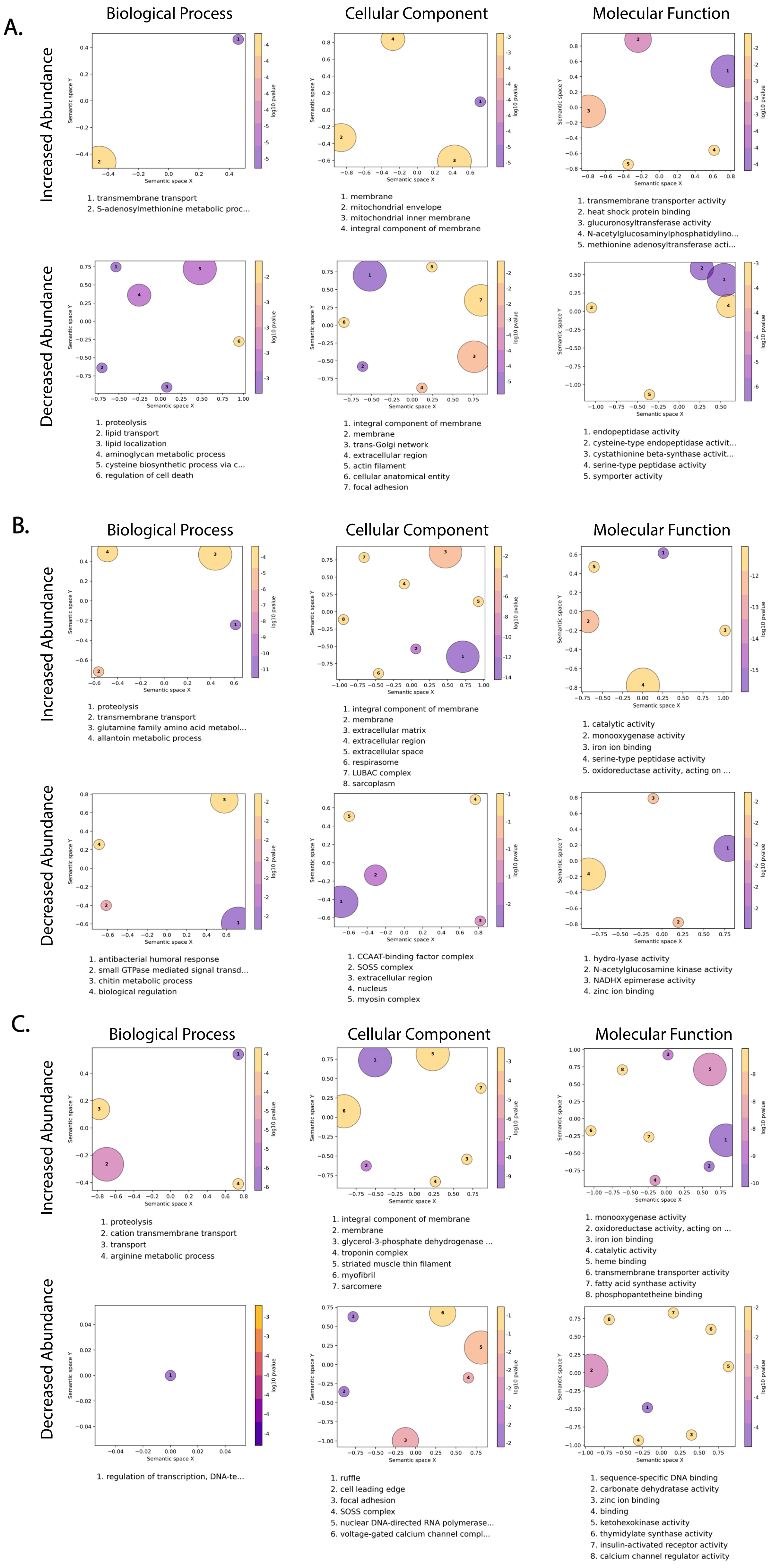
A.
Biological Process
Cellular Component
Molecular Function
Increased Abundance
Decreased Abundance
Semantic space X
Semantic space Y
log10 pvalue
1. transmembrane transport
2. S-adenosylmethionine metabolic proc...
1. membrane
2. mitochondrial envelope
3. mitochondrial inner membrane
4. integral component of membrane
1. transmembrane transporter activity
2. heat shock protein binding
3. glucuronosyltransferase activity
4. N-acetylglucosaminylphosphatidylino...
5. methionine adenosyltransferase acti...
1. proteolysis
2. lipid transport
3. lipid localization
4. aminoglycan metabolic process
5. cysteine biosynthetic process via c...
6. regulation of cell death
1. integral component of membrane
2. membrane
3. trans-Golgi network
4. extracellular region
5. actin filament
6. cellular anatomical entity
7. focal adhesion
1. endopeptidase activity
2. cysteine-type endopeptidase activit...
3. cystathionine beta-synthase activit...
4. serine-type peptidase activity
5. symporter activity
B.
Biological Process
Cellular Component
Molecular Function
Increased Abundance
Decreased Abundance
1. proteolysis
2. transmembrane transport
3. glutamine family amino acid metabol...
4. allantoin metabolic process
1. integral component of membrane
2. membrane
3. extracellular matrix
4. extracellular region
5. extracellular space
6. respirasome
7. LUBAC complex
8. sarcoplasm
1. catalytic activity
2. monooxygenase activity
3. iron ion binding
4. serine-type peptidase activity
5. oxidoreductase activity, acting on ...
1. antibacterial humoral response
2. small GTPase mediated signal transd...
3. chitin metabolic process
4. biological regulation
1. CCAAT-binding factor complex
2. SOSS complex
3. extracellular region
4. nucleus
5. myosin complex
1. hydro-lyase activity
2. N-acetylglucosamine kinase activity
3. NADHX epimerase activity
4. zinc ion binding
C.
Biological Process
Cellular Component
Molecular Function
Increased Abundance
Decreased Abundance
1. proteolysis
2. cation transmembrane transport
3. transport
4. arginine metabolic process
1. integral component of membrane
2. membrane
3. glycerol-3-phosphate dehydrogenase ...
4. troponin complex
5. striated muscle thin filament
6. myofibril
7. sarcomere
1. monooxygenase activity
2. oxidoreductase activity, acting on ...
3. iron ion binding
4. catalytic activity
5. heme binding
6. transmembrane transporter activity
7. fatty acid synthase activity
8. phosphopantetheine binding
1. regulation of transcription, DNA-te...
1. ruffle
2. cell leading edge
3. focal adhesion
4. SOSS complex
5. nuclear DNA-directed RNA polymerase...
6. voltage-gated calcium channel compl...
1. sequence-specific DNA binding
2. carbonate dehydratase activity
3. zinc ion binding
4. binding
5. ketohexokinase activity
6. thymidylate synthase activity
7. insulin-activated receptor activity
8. calcium channel regulator activity

**Supplemental Figure 2: Gene Ontology analysis of COL.*w*Mel transcripts post blood-feeding.** GO terms associated with the differentially expressed transcripts in COL.*w*Mel midguts at 7dpf **(A)** and carcasses at 4 and 7dpf **(B,C)**. The top 10 GO terms from each category (Biological Process, Cellular Component, Molecular Function), determined by topGO, were run in the GO Figure! pipeline to combine semantically similar terms and reduce redundancy. Terms are ranked by lowest log10(p-value). The size of each graphical point corresponds to the number of topGO terms associated with the listed summarizing term.

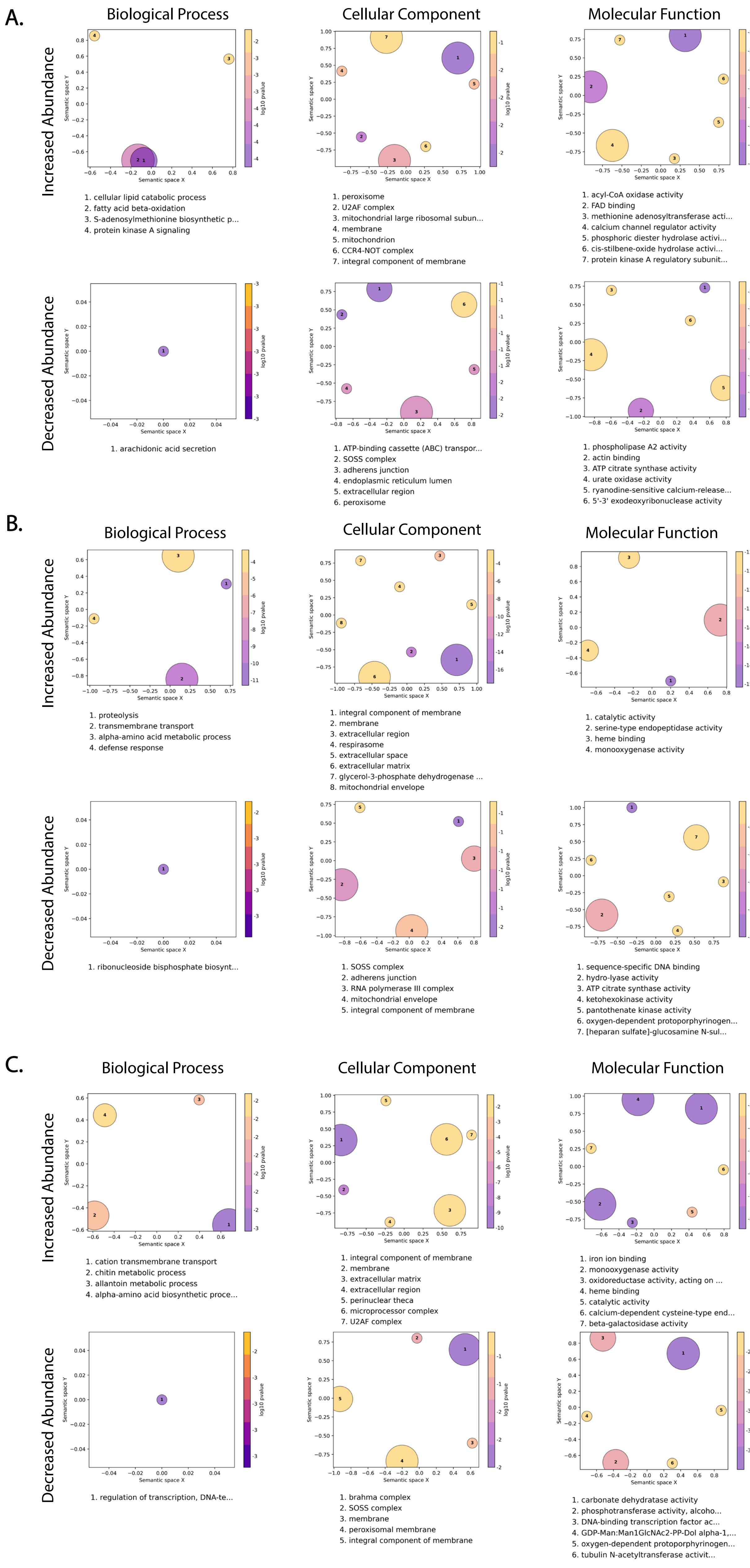
A.
Biological Process
Cellular Component
Molecular Function
Increased Abundance
Decreased Abundance
Semantic space X
Semantic space Y
log10 pvalue
1. cellular lipid catabolic process
2. fatty acid beta-oxidation
3. S-adenosylmethionine biosynthetic p...
4. protein kinase A signaling
1. peroxisome
2. U2AF complex
3. mitochondrial large ribosomal subun...
4. membrane
5. mitochondrion
6. CCR4-NOT complex
7. integral component of membrane
1. acyl-CoA oxidase activity
2. FAD binding
3. methionine adenosyltransferase acti...
4. calcium channel regulator activity
5. phosphoric diester hydrolase activi...
6. cis-stilbene-oxide hydrolase activi...
7. protein kinase A regulatory subunit...
1. arachidonic acid secretion
1. ATP-binding cassette (ABC) transpor...
2. SOSS complex
3. adherens junction
4. endoplasmic reticulum lumen
5. extracellular region
6. peroxisome
1. phospholipase A2 activity
2. actin binding
3. ATP citrate synthase activity
4. urate oxidase activity
5. ryanodine-sensitive calcium-release...
6. 5'-3' exodeoxyribonuclease activity
B.
Biological Process
Cellular Component
Molecular Function
Increased Abundance
Decreased Abundance
1. proteolysis
2. transmembrane transport
3. alpha-amino acid metabolic process
4. defense response
1. integral component of membrane
2. membrane
3. extracellular region
4. respirasome
5. extracellular space
6. extracellular matrix
7. glycerol-3-phosphate dehydrogenase ...
8. mitochondrial envelope
1. catalytic activity
2. serine-type endopeptidase activity
3. heme binding
4. monooxygenase activity
1. ribonucleoside bisphosphate biosynt...
1. SOSS complex
2. adherens junction
3. RNA polymerase III complex
4. mitochondrial envelope
5. integral component of membrane
1. sequence-specific DNA binding
2. hydro-lyase activity
3. ATP citrate synthase activity
4. ketohexokinase activity
5. pantothenate kinase activity
6. oxygen-dependent protoporphyrinogen...
7. [heparan sulfate]-glucosamine N-sul...
C.
Biological Process
Cellular Component
Molecular Function
Increased Abundance
Decreased Abundance
1. cation transmembrane transport
2. chitin metabolic process
3. allantoin metabolic process
4. alpha-amino acid biosynthetic proce...
1. integral component of membrane
2. membrane
3. extracellular matrix
4. extracellular region
5. perinuclear theca
6. microprocessor complex
7. U2AF complex
1. iron ion binding
2. monooxygenase activity
3. oxidoreductase activity, acting on ...
4. heme binding
5. catalytic activity
6. calcium-dependent cysteine-type end...
7. beta-galactosidase activity
1. regulation of transcription, DNA-te...
1. brahma complex
2. SOSS complex
3. membrane
4. peroxisomal membrane
5. integral component of membrane
1. carbonate dehydratase activity
2. phosphotransferase activity, alcoho...
3. DNA-binding transcription factor ac...
4. GDP-Man:Man1GlcNAc2-PP-Dol alpha-1,...
5. oxygen-dependent protoporphyrinogen...
6. tubulin N-acetyltransferase activit...

**Supplemental Figure 3: Gene Ontology analysis of COL.*w*Mel transcripts post blood-feeding on ZIKV-infected mice.** GO terms associated with the differentially expressed transcripts in COL.*w*Mel midguts at 7dpf **(A)** and carcasses at 4 and 7dpf **(B,C)** on a ZIKV-infected bloodmeal. The top 10 GO terms from each category (Biological Process, Cellular Component, Molecular Function), determined by topGO, were run in the GO Figure! pipeline to combine semantically similar terms and reduce redundancy. Terms are ranked by lowest log10(p-value). The size of each graphical point corresponds to the number of topGO terms associated with the listed summarizing term.

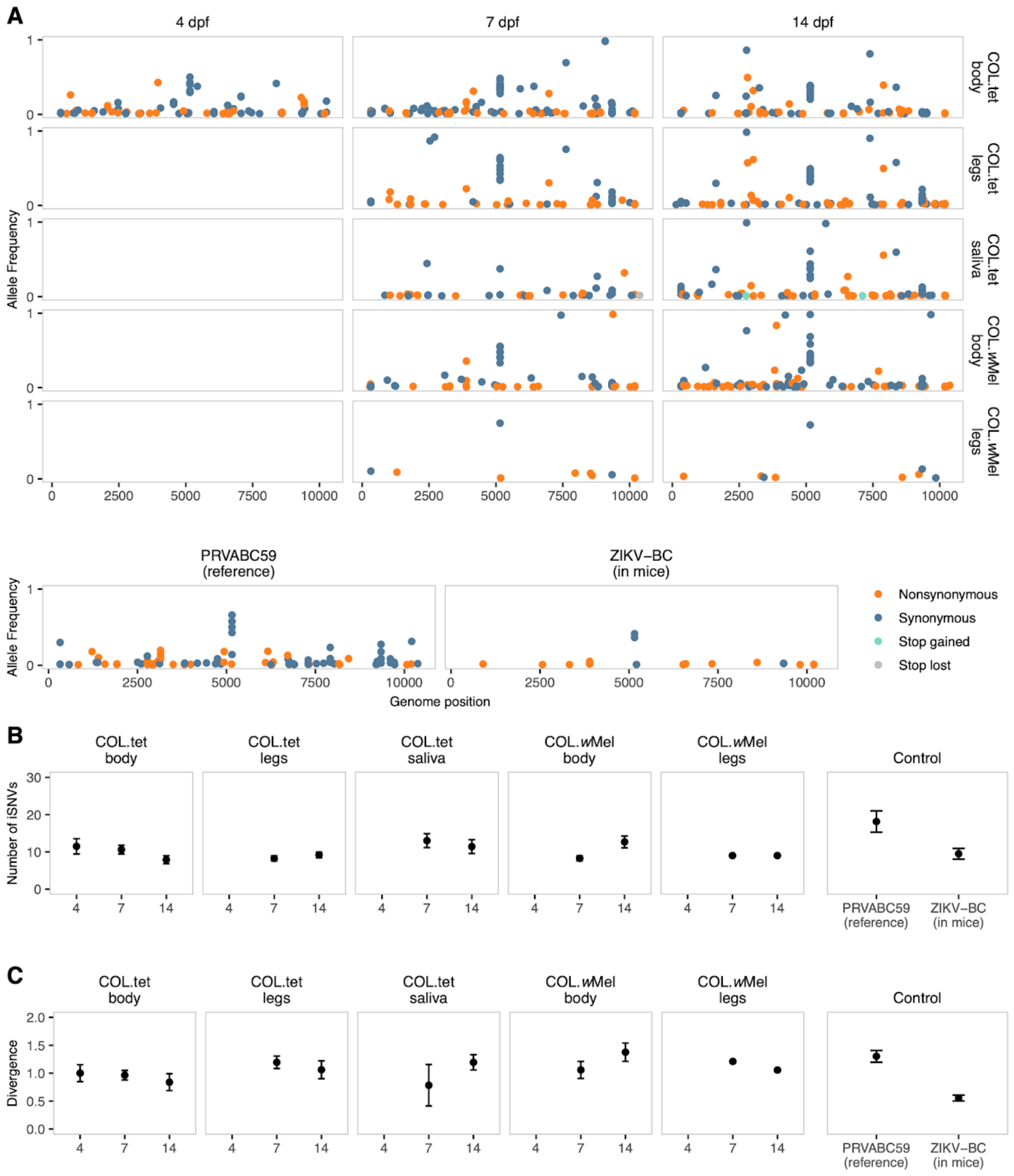


**Supplemental Figure 4: iSNVs occur across the ZIKV genome at similar levels. (A)**. iSNVs ≥ 1% are plotted along the PRVABC59 genome and colored by mutation type: nonsynonymous (orange), synonymous (blue), stop gained (green), and stop lost (grey). **(B)**. Number of iSNVs per sample are plotted across groups and time points. **(C)**. The per-sample divergence (total allele frequency) is plotted as in B. All groups in B and C underwent 10,000 Bayesian bootstrap replicates, from which mean values and standard deviations were calculated and plotted.

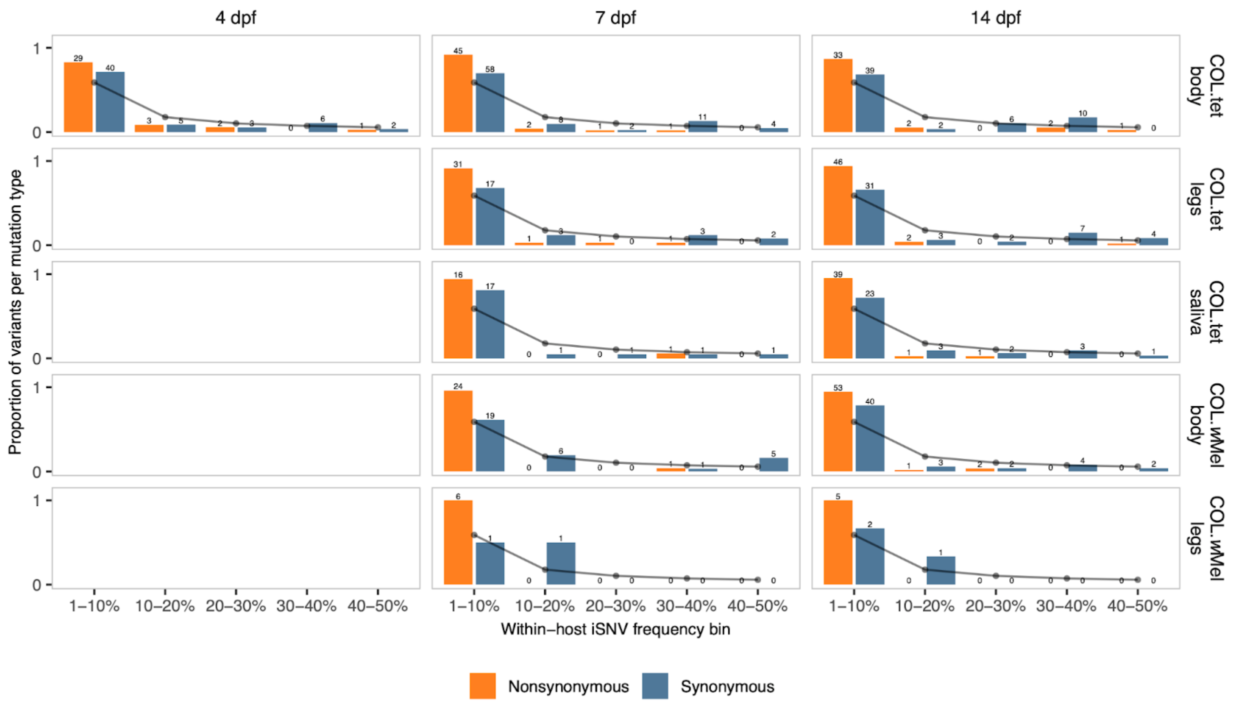


**Supplemental Figure 5: iSNV frequency-distribution spectra.** The proportion of variants per mutation type was calculated for nonsynonymous (orange) and synonymous (blue) mutation types. For each group and mutation type, the number of mutations that fell within a within-host iSNV frequency bin was divided by the total number of mutations (≤ 50% allele frequency). The grey dots and connecting lines denote the neutral expectation proportion for each frequency bin, assuming neutral selection and constant population size, modeled as following an inverse distribution.

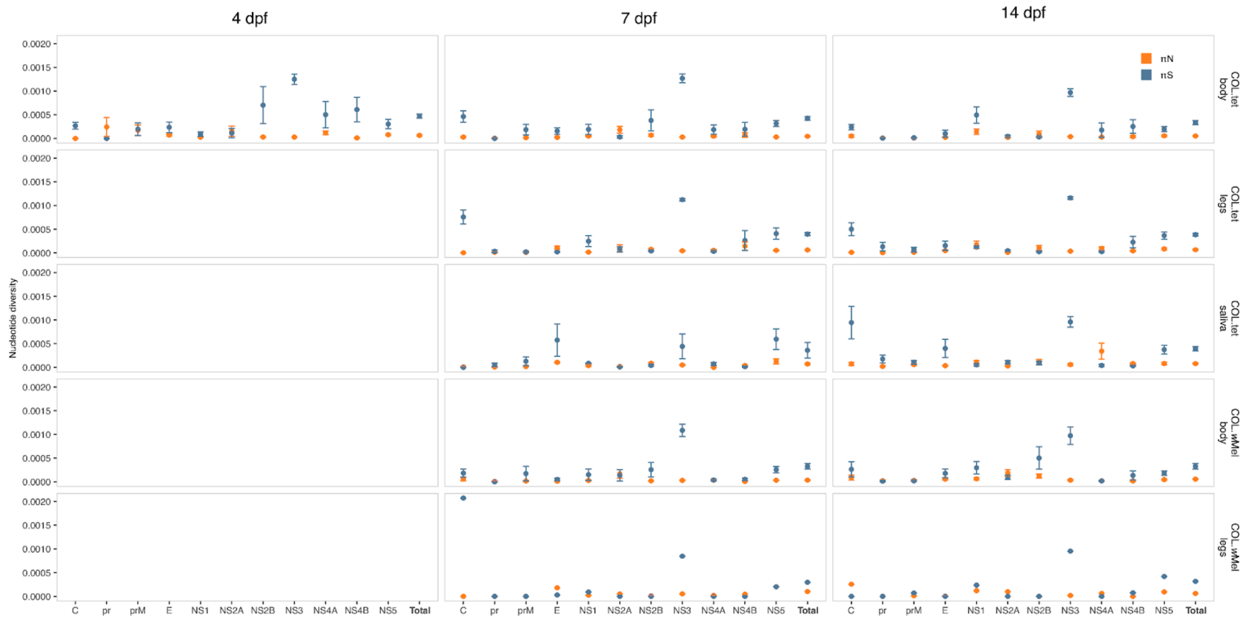


**Supplemental Figure 6: Gene-wise nucleotide diversity.** Per-gene nucleotide diversity is quantified for nonsynonymous (πN; orange) and synonymous (πS; blue) sites across all ZIKV plaque-positive tissues collected from COL.*w*Mel and COL.tet mosquitoes. All groups underwent 10,000 Bayesian bootstrap replicates, from which mean values and standard deviations were calculated and plotted.

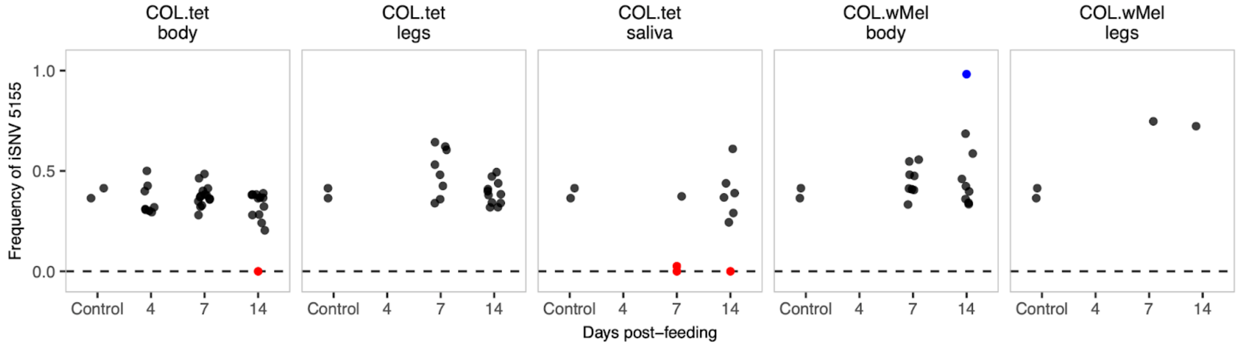


**Supplemental Figure 7: iSNV 5155 persists at intermediate levels in all groups.** The allele frequency of iSNV 5155 is plotted over time in all experimental groups. Samples were colored by allele frequencies: <5% (red), 5–95% (black), >95% (blue). If iSNV 5155 was not detected in a sample, it was assigned the allele frequency 0.
