## Supplemental Tables for "*Wolbachia*-mediated resistance to Zika virus infection in *Aedes aegypti* is dominated by diverse transcriptional regulation and weak evolutionary pressures"

| Table S1. Mosquito innate immune genes differentially expressed in COL.wMel relative to COL.tet. | | | | | | |
|---|---|---|---|---|---|---|
| **Vectorbase ID** | **Gene Name** | **Product Description** | **GO term: CC** | **GO term: MF** | **GO term: BP** | **Sample Group** |
| AAEL000037 | CLIPB35 | Clip-Domain Serine Protease family B. | extracellular region | hydrolase activity;peptidase activity;serine-type endopeptidase activity;serine-type peptidase activity | proteolysis | carcass_4, carcass_7 |
| AAEL000227 | SCRB8 | Class B Scavenger Receptor (CD36 domain). | integral component of membrane;membran e | N/A | N/A | carcass_4 |
| AAEL000234 | SCRB7 | Class B Scavenger Receptor (CD36 domain). | membrane | N/A | N/A | carcass_4 |
| AAEL000652 | GNBPA2 | Gram-Negative Binding Protein (GNBP) or Beta-1 3-Glucan Binding Protein (BGBP). | N/A | carbohydrate binding;hydrolase activity, hydrolyzing O-glycosyl compounds | carbohydrate metabolic process | midgut_7 |
| AAEL000726 | N/A | fibrinogen and fibronectin | N/A | N/A | N/A | carcass_7 |
| AAEL000749 | N/A | Angiopoietin-like protein variant (Fragment) [Source:UniProtKB/Tr EMBL;Acc:Q1HRV2] | N/A | N/A | N/A | carcass_7 |
| AAEL000760 | CLIPB30 | Clip-Domain Serine Protease family B. | extracellular region | hydrolase activity;peptidase activity;serine-type endopeptidase activity;serine-type peptidase activity | proteolysis | carcass_7 |
| AAEL001401 | LRIM10A | leucine-rich immune protein (Short) | N/A | protein binding | N/A | carcass_4, carcass_7 |
| AAEL001402 | LRIM10B | leucine-rich immune protein (Short) | N/A | protein binding | N/A | carcass_4, carcass_7 |
| AAEL001414 | LRIM9 | leucine-rich immune protein (Short) | N/A | protein binding | N/A | carcass_4, carcass_4, carcass_7 |
| AAEL001417 | LRIM7 | leucine-rich immune protein (Short) | N/A | protein binding | N/A | carcass_4, carcass_7 |
| AAEL001420 | LRIM8 | leucine-rich immune protein (Short) | N/A | protein binding | N/A | carcass_4, carcass_7 |
| AAEL001650 | N/A | ML domain-containing protein [Source:UniProtKB/Tr EMBL;Acc:A0A1S4EZ F1] | N/A | N/A | N/A | midgut_7 |
| AAEL001794 | N/A | macroglobulin/compl ement | extracellular region;extracellular space | endopeptidase inhibitor activity | negative regulation of endopeptidase activity | carcass_4, carcass_7 |
| AAEL002309 | TPX4 | Thioredoxin Peroxidase. | obsolete cell | antioxidant activity;oxidoreductas | cell redox homeostasis;cellular | midgut_7 |

| | | | | | | |
|---|---|---|---|---|---|---|
| | | | | e activity;peroxiredoxin activity | oxidant detoxification;obsolete oxidation-reduction process | |
| AAEL002601 | CLIPA1 | Clip-Domain Serine Protease family A. Protease homologue. | N/A | serine-type endopeptidase activity | proteolysis | carcass_4 |
| AAEL002720 | SRPN20 | Serine Protease Inhibitor (serpin) likely cleavage at V/V. | extracellular space | N/A | N/A | carcass_4 |
| AAEL002731 | SRPN14 | Serine Protease Inhibitor (serpin) homologue - unlikely to be inhibitory. | extracellular space | N/A | N/A | carcass_4, carcass_7 |
| AAEL003156 | N/A | fibrinogen and fibronectin | N/A | N/A | N/A | carcass_4 |
| AAEL003182 | SRPN26 | Serine Protease Inhibitor (serpin) homologue - unlikely to be inhibitory. | extracellular space | N/A | N/A | midgut_4 |
| AAEL003253 | CLIPB13B | Clip-Domain Serine Protease family B. | N/A | serine-type endopeptidase activity | proteolysis | carcass_4, carcass_7 |
| AAEL003294 | N/A | fibrinogen and fibronectin | N/A | N/A | N/A | midgut_7 |
| AAEL003389 | ATT | attacin anti-microbial peptide | extracellular region;extracellular space | N/A | antibacterial humoral response;defense response to bacterium | carcass_4 |
| AAEL003439 | CASPS18 | caspase (short) | N/A | cysteine-type endopeptidase activity;cysteine-type peptidase activity | proteolysis | midgut_7 |
| AAEL003444 | CASPS19 | caspase (short) | N/A | cysteine-type endopeptidase activity;cysteine-type peptidase activity;hydrolase activity;peptidase activity | proteolysis | midgut_7 |
| AAEL003631 | CLIPB41 | Clip-Domain Serine Protease family B. | N/A | hydrolase activity;peptidase activity;serine-type endopeptidase activity;serine-type peptidase activity | proteolysis | carcass_4 |
| AAEL003697 | SRPN17 | Serine Protease Inhibitor (serpin) homologue - unlikely to be inhibitory. | extracellular space | N/A | N/A | carcass_4, carcass_7 |
| AAEL003712 | LYSC10 | C-Type Lysozyme (Lys-E). | N/A | lysozyme activity | N/A | midgut_7 |
| AAEL003723 | LYSC11 | C-Type Lysozyme (Lys-A). | N/A | lysozyme activity | N/A | midgut_4 |
| AAEL004390 | HPX8B | heme peroxidase | N/A | heme binding;peroxidase activity | cellular oxidant detoxification;obsolete oxidation-reduction process;oogenesis;response to oxidative stress | midgut_4 |

| | | | | | | |
|---|---|---|---|---|---|---|
| AAEL004833 | N/A | unspecified product | extracellular region | N/A | defense response to bacterium | carcass_4, carcass_4 |
| AAEL004978 | N/A | DEAD box ATP-dependent RNA helicase | N/A | ATP binding;helicase activity;hydrolase activity;nucleic acid binding;nucleotide binding | N/A | midgut_4 |
| AAEL004979 | CLIPD2 | Clip-Domain Serine Protease family D. | N/A | hydrolase activity;peptidase activity;serine-type endopeptidase activity;serine-type peptidase activity | proteolysis | carcass_4 |
| AAEL005108 | MNSOD2 | manganese-iron (Mn-Fe) superoxide dismutase | N/A | metal ion binding;oxidoreductase activity;superoxide dismutase activity | obsolete oxidation-reduction process;removal of superoxide radicals;superoxide metabolic process | carcass_4, carcass_7 |
| AAEL005293 | GALE8A | Galectin [Source:UniProtKB/TrEMBL;Acc:Q16ND5] | N/A | carbohydrate binding | N/A | midgut_7 |
| AAEL005482 | CTL18 | C-Type Lectin (CTL). | N/A | carbohydrate binding | N/A | midgut_4 |
| AAEL005641 | CTLGA5 | C-Type Lectin (CTL) - galactose binding. | N/A | N/A | N/A | midgut_7 |
| AAEL005956 | CASPS16 | caspase (short) | N/A | cysteine-type endopeptidase activity;cysteine-type peptidase activity | proteolysis | midgut_4, carcass_4 |
| AAEL006161 | CLIPB31 | Clip-Domain Serine Protease family B | extracellular region | hydrolase activity;peptidase activity;serine-type endopeptidase activity;serine-type peptidase activity | proteolysis | carcass_4 |
| AAEL006355 | SCRC1 | Class C Scavenger Receptor (Sushi/SCR/CCP MAM and Somatomedin B domains). | integral component of membrane;membrane | N/A | N/A | midgut_7 |
| AAEL006361 | SCRC2 | Class C Scavenger Receptor (Sushi/SCR/CCP MAM and Somatomedin B domains). | integral component of membrane;membrane | N/A | N/A | midgut_4, midgut_7 |
| AAEL006377 | LRIM31 | leucine-rich immune protein (Coil-less) | N/A | protein binding | N/A | carcass_4 |
| AAEL006674 | CLIPB29 | Clip-Domain Serine Protease family B. | extracellular region | hydrolase activity;peptidase activity;serine-type endopeptidase activity;serine-type peptidase activity | proteolysis | carcass_4 |
| AAEL006704 | N/A | fibrinogen and fibronectin | N/A | N/A | N/A | midgut_7 |
| AAEL006854 | N/A | Niemann-Pick Type C-2, putative | N/A | N/A | intracellular cholesterol transport | midgut_4, midgut_7 |
| AAEL007103 | LRIM15 | leucine-rich immune protein (TM) | integral component of | protein binding | N/A | carcass_4 |

| | | | | | | |
|---|---|---|---|---|---|---|
| | | | membrane;membrane | | | |
| AAEL007224 | LRIM22 | leucine-rich immune protein (Coil-less) | N/A | protein binding | N/A | carcass_4, carcass_7 |
| AAEL007624 | REL2 | IMD pathway signalling NF-kappaB Relish-like transcription factor | cytoplasm;host cell nucleus;nucleus | DNA binding;DNA-binding transcription factor activity;protein binding | regulation of transcription, DNA-templated | midgut_7 |
| AAEL007696 | REL1A | TOLL pathway signalling NF-kappaB Relish-like transcription factor | cytoplasm;host cell nucleus;nucleus | DNA binding;DNA-binding transcription factor activity | regulation of transcription, DNA-templated | carcass_4, carcass_7 |
| AAEL007823 | N/A | PIWI | N/A | nucleic acid binding;protein binding | gene silencing by RNA | midgut_7 |
| AAEL007993 | CLIPB27 | Clip-Domain Serine Protease family B. | extracellular region | hydrolase activity;peptidase activity;serine-type endopeptidase activity;serine-type peptidase activity | proteolysis | midgut_4, midgut_7 |
| AAEL008098 | PIWI2 | PIWI | N/A | nucleic acid binding;protein binding | gene silencing by RNA | midgut_4 |
| AAEL008370 | SCRB17 | Class B Scavenger Receptor (CD36 domain). | integral component of membrane;membrane | N/A | N/A | carcass_4, carcass_7 |
| AAEL008658 | LRIM16 | leucine-rich immune protein (TM) | integral component of membrane;membrane | protein binding | N/A | midgut_4, midgut_7 |
| AAEL008668 | CLIPB22 | Clip-Domain Serine Protease family B. | N/A | serine-type endopeptidase activity | proteolysis | carcass_4 |
| AAEL009178 | GNBPB4 | Gram-Negative Binding Protein (GNBP) or Beta-1 3-Glucan Binding Protein (BGBP). | N/A | hydrolase activity, hydrolyzing O-glycosyl compounds | carbohydrate metabolic process | carcass_7 |
| AAEL009384 | N/A | fibrinogen and fibronectin | N/A | N/A | N/A | midgut_7 |
| AAEL009474 | PGRPS1 | Peptidoglycan Recognition Protein (Short) | N/A | N-acetylmuramoyl-L-alanine amidase activity;peptidoglycan binding;zinc ion binding | immune system process;innate immune response;peptidoglycan catabolic process | carcass_4 |
| AAEL009556 | N/A | Niemann-Pick Type C-2, putative | N/A | N/A | intracellular cholesterol transport | midgut_4, midgut_7 |
| AAEL009792 | LRIM25 | leucine-rich immune protein (Coil-less) | N/A | protein binding | N/A | carcass_7 |
| AAEL009842 | GALE12 | Galectin [Source:UniProtKB/TrEMBL;Acc:Q16UP1] | N/A | N/A | N/A | midgut_4, midgut_7 |
| AAEL009850 | GALE14 | Galectin [Source:UniProtKB/TrEMBL;Acc:Q16UP0] | N/A | carbohydrate binding | N/A | midgut_4, midgut_7 |

| | | | | | | |
|---|---|---|---|---|---|---|
| AAEL009894 | LRIM21 | leucine-rich immune protein (Coil-less) | integral component of membrane;membrane | protein binding | N/A | midgut_4, midgut_7 |
| AAEL010125 | LRIM17 | leucine-rich immune protein (Coil-less) | N/A | protein binding | N/A | carcass_4 |
| AAEL010128 | LRIM4 | leucine-rich immune protein (Long) | N/A | protein binding | N/A | carcass_4, carcass_7 |
| AAEL010132 | LRIM3 | leucine-rich immune protein (Long) | N/A | protein binding | N/A | carcass_4 |
| AAEL010769 | SRPN6 | Serine Protease Inhibitor (serpin) likely cleavage at S/A. | extracellular space | N/A | N/A | carcass_7 |
| AAEL011009 | N/A | fibrinogen and fibronectin | N/A | N/A | N/A | carcass_4, carcass_7 |
| AAEL011407 | CTL20 | C-Type Lectin (CTL20) | N/A | N/A | N/A | midgut_4 |
| AAEL011408 | CTL21 | C-Type Lectin (CTL21) | N/A | N/A | N/A | carcass_4, carcass_7 |
| AAEL011453 | CTL14 | C-Type Lectin (CTL14) | N/A | N/A | N/A | carcass_4, carcass_7 |
| AAEL011455 | CTLMA12 | C-Type Lectin (CTLMA12) - mannose binding | N/A | carbohydrate binding | N/A | carcass_4 |
| AAEL011621 | CTLMA13 | C-Type Lectin (CTL) - mannose binding. | N/A | carbohydrate binding;serine-type endopeptidase activity | proteolysis | carcass_4, carcass_7 |
| AAEL011633 | N/A | fibrinogen and fibronectin | N/A | N/A | N/A | carcass_4 |
| AAEL011763 | PPO3 | prophenoloxidase | N/A | metal ion binding;oxidoreductase activity | obsolete oxidation-reduction process | carcass_4 |
| AAEL011764 | PPO10 | prophenoloxidase | N/A | metal ion binding;oxidoreductase activity | obsolete oxidation-reduction process | carcass_4, carcass_7 |
| AAEL012003 | GALE6B | Galectin [Source:UniProtKB/TrEMBL;Acc:Q17AG2] | N/A | carbohydrate binding | N/A | midgut_7 |
| AAEL012086 | LRIM1 | leucine-rich immune protein (Long) | N/A | protein binding | N/A | carcass_4, carcass_7 |
| AAEL012255 | LRIM13 | leucine-rich immune protein (Short) | N/A | protein binding | N/A | carcass_4, carcass_7 |
| AAEL012380 | PGRPLA | Peptidoglycan Recognition Protein (Long) | integral component of membrane;membrane | N-acetylmuramoyl-L-alanine amidase activity;zinc ion binding | peptidoglycan catabolic process | midgut_4, midgut_7 |
| AAEL012471 | DOME | JAKSTAT pathway signalling Transmembrane Receptor Domeless. | integral component of membrane;membrane | cytokine receptor activity;protein binding | cytokine-mediated signaling pathway | midgut_7 |
| AAEL012763 | LRIM24 | leucine-rich immune protein (Coil-less) | N/A | protein binding | N/A | carcass_4 |

| | | | | | | |
|---|---|---|---|---|---|---|
| AAEL012767 | LRIM5 | leucine-rich immune protein (Short) | N/A | protein binding | N/A | carcass_4 |
| AAEL012911 | LRIM18 | leucine-rich immune protein (Coil-less) | integral component of membrane;membrane | protein binding | N/A | carcass_4 |
| AAEL013245 | CLIPB28 | Clip-Domain Serine Protease family B. | N/A | peptidase activity;serine-type endopeptidase activity | proteolysis | carcass_7, carcass_7 |
| AAEL013417 | N/A | fibrinogen and fibronectin | N/A | N/A | N/A | carcass_4 |
| AAEL013496 | PPO8 | prophenoloxidase | N/A | metal ion binding;oxidoreductase activity | obsolete oxidation-reduction process | carcass_4 |
| AAEL013692 | PIWI3 | PIWI | N/A | nucleic acid binding;protein binding | gene silencing by RNA | midgut_4 |
| AAEL014348 | CASPS8 | caspase (short) | N/A | cysteine-type endopeptidase activity;cysteine-type peptidase activity;hydrolase activity;peptidase activity | proteolysis | midgut_7, midgut_7 |
| AAEL014382 | CTLMA14 | C-Type Lectin (CTL) - mannose binding. | N/A | carbohydrate binding | N/A | carcass_4 |
| AAEL014544 | PPO6 | prophenoloxidase | N/A | metal ion binding;oxidoreductase activity | obsolete oxidation-reduction process | carcass_4, carcass_7 |
| AAEL017536 | GRRP | holotricin glycine rich repeat protein (GRRP) anti-microbial peptide | N/A | N/A | N/A | carcass_4, carcass_4, carcass_4, carcass_7, carcass_7 |

CC, cellular component; MF, molecular function; BP, biological process.

| **Table S2. Mosquito innate immune genes differentially expressed in COL.wMel relative to COL.tet during ZIKV infection.** | | | | | | |
|---|---|---|---|---|---|---|
| **Vectorbase ID** | **Gene Name** | **Product Description** | **GO term: CC** | **GO term: MF** | **GO term: BP** | **Sample Group** |
| AAEL000037 | CLIPB35 | Clip-Domain Serine Protease family B. | extracellular region | hydrolase activity;peptidase activity;serine-type endopeptidase activity;serine-type peptidase activity | proteolysis | carcass_4 |
| AAEL000057 | TOLL5B | Toll-like receptor | integral component of membrane;membrane | protein binding | immune system process;innate immune response;signal transduction | carcass_4 |
| AAEL000227 | SCRB8 | Class B Scavenger Receptor (CD36 domain). | integral component of membrane;membrane | N/A | N/A | carcass_4 |
| AAEL000234 | SCRB7 | Class B Scavenger Receptor (CD36 domain). | membrane | N/A | N/A | carcass_7 |
| AAEL000652 | GNBPA2 | Gram-Negative Binding Protein (GNBP) or Beta-1 3-Glucan Binding Protein (BGBP). | N/A | carbohydrate binding;hydrolase activity, hydrolyzing O-glycosyl compounds | carbohydrate metabolic process | midgut_4 |
| AAEL000760 | CLIPB30 | Clip-Domain Serine Protease family B. | extracellular region | hydrolase activity;peptidase activity;serine-type endopeptidase activity;serine-type peptidase activity | proteolysis | carcass_4 |
| AAEL001077 | CLIPB45 | Clip-Domain Serine Protease family B. Protease homologue. | N/A | serine-type endopeptidase activity | proteolysis | carcass_4, carcass_7 |
| AAEL001401 | LRIM10A | leucine-rich immune protein (Short) | N/A | protein binding | N/A | carcass_4 |
| AAEL001402 | LRIM10B | leucine-rich immune protein (Short) | N/A | protein binding | N/A | carcass_4 |
| AAEL001414 | LRIM9 | leucine-rich immune protein (Short) | N/A | protein binding | N/A | carcass_4, carcass_4, carcass_7, carcass_7 |
| AAEL001417 | LRIM7 | leucine-rich immune protein (Short) | N/A | protein binding | N/A | carcass_4 |
| AAEL001420 | LRIM8 | leucine-rich immune protein (Short) | N/A | protein binding | N/A | carcass_4, carcass_7 |
| AAEL001794 | N/A | macroglobulin/complement | extracellular region;extracellular space | endopeptidase inhibitor activity | negative regulation of endopeptidase activity | carcass_4 |
| AAEL002126 | CLIPA15 | Clip-Domain Serine Protease family A. Protease homologue. | N/A | serine-type endopeptidase activity | proteolysis | carcass_4 |
| AAEL002601 | CLIPA1 | Clip-Domain Serine Protease family A. | N/A | serine-type endopeptidase | proteolysis | carcass_4 |

| | | | | | | |
|---|---|---|---|---|---|---|
| | | Protease homologue. | | activity | | |
| AAEL002720 | SRPN20 | Serine Protease Inhibitor (serpin) likely cleavage at V/V. | extracellular space | N/A | N/A | carcass_4 |
| AAEL002731 | SRPN14 | Serine Protease Inhibitor (serpin) homologue - unlikely to be inhibitory. | extracellular space | N/A | N/A | carcass_4 |
| AAEL003253 | CLIPB13B | Clip-Domain Serine Protease family B. | N/A | serine-type endopeptidase activity | proteolysis | carcass_4 |
| AAEL003631 | CLIPB41 | Clip-Domain Serine Protease family B. | N/A | hydrolase activity;peptidase activity;serine-type endopeptidase activity;serine-type peptidase activity | proteolysis | carcass_4, carcass_7 |
| AAEL003697 | SRPN17 | Serine Protease Inhibitor (serpin) homologue - unlikely to be inhibitory. | extracellular space | N/A | N/A | carcass_4 |
| AAEL003832 | DEFC | defensin anti-microbial peptide | extracellular region | N/A | defense response;defense response to bacterium;immune system process;innate immune response | carcass_4 |
| AAEL003841 | DEFA | Defensin-A [Source:UniProtKB/Swiss-Prot;Acc:P91793] | N/A | N/A | defense response | carcass_4 |
| AAEL003857 | N/A | INVERT_DEFENSINS domain-containing protein [Source:UniProtKB/TrEMBL;Acc:A0A1S4F676] | N/A | N/A | defense response | carcass_4 |
| AAEL004120 | N/A | Niemann-Pick Type C-2, putative | N/A | N/A | intracellular cholesterol transport | carcass_4 |
| AAEL004524 | CLIPC5B | Clip-Domain Serine Protease family C. | N/A | hydrolase activity;peptidase activity;serine-type endopeptidase activity;serine-type peptidase activity | proteolysis | carcass_4 |
| AAEL004833 | N/A | unspecified product | extracellular region | N/A | defense response to bacterium | carcass_4 |
| AAEL004979 | CLIPD2 | Clip-Domain Serine Protease family D. | N/A | hydrolase activity;peptidase activity;serine-type endopeptidase activity;serine-type peptidase activity | proteolysis | carcass_7 |
| AAEL005093 | CLIPB46 | Clip-Domain Serine Protease family B. | extracellular region | hydrolase activity;peptidase activity;serine-type endopeptidase activity;serine-type peptidase activity | proteolysis | carcass_4 |
| AAEL005108 | MNSOD2 | manganese-iron (Mn-Fe) superoxide dismutase | N/A | metal ion binding;oxidoreductase activity;superoxide | obsolete oxidation-reduction process;removal of | carcass_4, carcass_7 |

| | | | | | | |
|---|---|---|---|---|---|---|
| | | | | dismutase activity | superoxide radicals;superoxide metabolic process | |
| AAEL005431 | CLIPB37 | Clip-Domain Serine Protease family B. | N/A | hydrolase activity;peptidase activity;serine-type endopeptidase activity;serine-type peptidase activity | proteolysis | carcass_4 |
| AAEL005792 | CLIPE8 | Clip-Domain Serine Protease family E. Protease homologue. | N/A | peptidase activity;serine-type endopeptidase activity | proteolysis | carcass_4 |
| AAEL006161 | CLIPB31 | Clip-Domain Serine Protease family B | extracellular region | hydrolase activity;peptidase activity;serine-type endopeptidase activity;serine-type peptidase activity | proteolysis | carcass_7 |
| AAEL006377 | LRIM31 | leucine-rich immune protein (Coil-less) | N/A | protein binding | N/A | carcass_4, carcass_7 |
| AAEL006674 | CLIPB29 | Clip-Domain Serine Protease family B. | extracellular region | hydrolase activity;peptidase activity;serine-type endopeptidase activity;serine-type peptidase activity | proteolysis | carcass_4 |
| AAEL007224 | LRIM22 | leucine-rich immune protein (Coil-less) | N/A | protein binding | N/A | carcass_4, carcass_7 |
| AAEL007696 | REL1A | TOLL pathway signalling NF-kappaB Relish-like transcription factor | cytoplasm;host cell nucleus;nucleus | DNA binding;DNA-binding transcription factor activity | regulation of transcription, DNA-templated | carcass_4 |
| AAEL008370 | SCRB17 | Class B Scavenger Receptor (CD36 domain). | integral component of membrane;membran e | N/A | N/A | carcass_4 |
| AAEL009178 | GNBPB4 | Gram-Negative Binding Protein (GNBP) or Beta-1 3-Glucan Binding Protein (BGBP). | N/A | hydrolase activity, hydrolyzing O-glycosyl compounds | carbohydrate metabolic process | carcass_7 |
| AAEL009420 | SCRBQ1 | Class B Scavenger Receptor (CD36 domain). | integral component of membrane;membran e | N/A | N/A | carcass_7 |
| AAEL009474 | PGRPS1 | Peptidoglycan Recognition Protein (Short) | N/A | N-acetylmuramoyl-L-a lanine amidase activity;peptidoglycan binding;zinc ion binding | immune system process;innate immune response;peptidoglyc an catabolic process | carcass_4 |
| AAEL009842 | GALE12 | Galectin [Source:UniProtKB/Tr EMBL;Acc:Q16UP1] | N/A | N/A | N/A | carcass_4 |
| AAEL010128 | LRIM4 | leucine-rich immune protein (Long) | N/A | protein binding | N/A | carcass_4 |
| AAEL011009 | N/A | fibrinogen and fibronectin | N/A | N/A | N/A | carcass_4 |
| AAEL011407 | CTL20 | C-Type Lectin (CTL20) | N/A | N/A | N/A | carcass_7 |

| | | | | | | |
|---|---|---|---|---|---|---|
| AAEL011408 | CTL21 | C-Type Lectin (CTL21) | N/A | N/A | N/A | carcass_4, carcass_7 |
| AAEL011453 | CTL14 | C-Type Lectin (CTL14) | N/A | N/A | N/A | carcass_4, carcass_7 |
| AAEL011455 | CTLMA12 | C-Type Lectin (CTLMA12) - mannose binding | N/A | carbohydrate binding | N/A | carcass_4 |
| AAEL011621 | CTLMA13 | C-Type Lectin (CTL) - mannose binding. | N/A | carbohydrate binding;serine-type endopeptidase activity | proteolysis | carcass_4, carcass_7 |
| AAEL011633 | N/A | fibrinogen and fibronectin | N/A | N/A | N/A | carcass_4, carcass_7 |
| AAEL011634 | N/A | fibrinogen and fibronectin | N/A | N/A | N/A | carcass_7 |
| AAEL011763 | PPO3 | prophenoloxidase | N/A | metal ion binding;oxidoreductase activity | obsolete oxidation-reduction process | carcass_4, carcass_7 |
| AAEL011764 | PPO10 | prophenoloxidase | N/A | metal ion binding;oxidoreductase activity | obsolete oxidation-reduction process | carcass_4, carcass_7 |
| AAEL012086 | LRIM1 | leucine-rich immune protein (Long) | N/A | protein binding | N/A | carcass_4 |
| AAEL012135 | GALE2 | Galectin [Source:UniProtKB/TrEMBL;Acc:Q16MZ7] | N/A | carbohydrate binding | N/A | carcass_4 |
| AAEL012255 | LRIM13 | leucine-rich immune protein (Short) | N/A | protein binding | N/A | midgut_7, carcass_4, carcass_7 |
| AAEL012410 | AGO1b | eukaryotic translation initiation factor 2C | N/A | nucleic acid binding;protein binding | N/A | midgut_4 |
| AAEL012538 | LRIM6 | leucine-rich immune protein (Short) | N/A | protein binding | N/A | carcass_4, carcass_7 |
| AAEL012767 | LRIM5 | leucine-rich immune protein (Short) | N/A | protein binding | N/A | carcass_4 |
| AAEL013245 | CLIPB28 | Clip-Domain Serine Protease family B. | N/A | peptidase activity;serine-type endopeptidase activity | proteolysis | carcass_4, carcass_4 |
| AAEL013417 | N/A | fibrinogen and fibronectin | N/A | N/A | N/A | carcass_7 |
| AAEL013434 | N/A | spaetzle-like cytokine | N/A | N/A | N/A | carcass_7 |
| AAEL013496 | PPO8 | prophenoloxidase | N/A | metal ion binding;oxidoreductase activity | obsolete oxidation-reduction process | carcass_4, carcass_7 |
| AAEL013501 | PPO4 | prophenoloxidase | N/A | metal ion binding;oxidoreductase activity | obsolete oxidation-reduction process | carcass_4 |
| AAEL014349 | CLIPB15 | Clip-Domain Serine Protease family B. | extracellular region | hydrolase activity;peptidase activity;serine-type endopeptidase activity;serine-type peptidase activity | proteolysis | carcass_4 |

| | | | | | | |
|---|---|---|---|---|---|---|
| AAEL014382 | CTLMA14 | C-Type Lectin (CTL) - mannose binding. | N/A | carbohydrate binding | N/A | carcass_4 |
| AAEL017249 | SRPN24 | Serine Protease Inhibitor (serpin) homologue - unlikely to be inhibitory. | extracellular space | N/A | N/A | carcass_4 |
| AAEL017536 | GRRP | holotricin glycine rich repeat protein (GRRP) anti-microbial peptide | N/A | N/A | N/A | carcass_4, carcass_4, carcass_4, carcass_7, carcass_7 |

CC, cellular component; MF, molecular function; BP, biological process.

| Table S3. Genes differentially expressed in ZIKV-exposed COL.*w*Mel carcasses 7dpf | | |
|---|---|---|
| **Gene ID** | **Product Description** | **Gene Name or Symbol** |
| **AAEL000102** | unspecified product | N/A |
| **AAEL000311** | unspecified product | N/A |
| **AAEL000415** | AMP dependent coa ligase | N/A |
| **AAEL000566** | unspecified product | N/A |
| **AAEL000658** | unspecified product | N/A |
| **AAEL000859** | unspecified product | N/A |
| **AAEL001062** | unspecified product | N/A |
| **AAEL001091** | Malic enzyme [Source:UniProtKB/TrEMBL;Acc:A0A1S4EXR8] | N/A |
| **AAEL001209** | sodium-dependent phosphate transporter | N/A |
| **AAEL001232** | tubulointerstitial nephritis antigen | N/A |
| **AAEL001293** | unspecified product | N/A |
| **AAEL001307** | SEC14, putative | N/A |
| **AAEL001818** | unspecified product | N/A |
| **AAEL002378** | Carboxylic ester hydrolase [Source:UniProtKB/TrEMBL;Acc:A0A0P6IYI7] | N/A |
| **AAEL002416** | short-chain dehydrogenase | N/A |
| **AAEL002467** | unspecified product | N/A |
| **AAEL002796** | l-asparaginase i | N/A |
| **AAEL002978** | leucyl aminopeptidase, putative | N/A |
| **AAEL003002** | unspecified product | N/A |
| **AAEL003803** | unspecified product | N/A |
| **AAEL004297** | ATP-citrate synthase | N/A |
| **AAEL004941** | cytochrome P450 | CYP6AK1 |
| **AAEL004974** | beta-1,3-glucuronyltransferase s, p | N/A |
| **AAEL005147** | unspecified product | N/A |
| **AAEL005199** | Carboxylic ester hydrolase [Source:UniProtKB/TrEMBL;Acc:A0A1S4F9U8] | N/A |
| **AAEL005256** | unspecified product | N/A |
| **AAEL005293** | Galectin [Source:UniProtKB/TrEMBL;Acc:Q16ND5] | GALE8A |
| **AAEL005428** | unspecified product | N/A |
| **AAEL005515** | heterogeneous nuclear ribonucleoprotein | N/A |
| **AAEL005790** | malic enzyme | N/A |
| **AAEL005992** | adam (a disintegrin and metalloprotease) | N/A |
| **AAEL006721** | 2-oxoglutarate dehydrogenase | N/A |
| **AAEL006883** | unspecified product | N/A |
| **AAEL007010** | cytochrome P450 | CYP6AG4 |
| **AAEL007029** | tropomodulin | N/A |
| **AAEL007271** | basic helix-loop-helix zip transcription factor | N/A |
| **AAEL007381** | unspecified product | N/A |
| **AAEL007653** | allantoinase | N/A |
| **AAEL007914** | discs large protein | N/A |

| | | |
|---|---|---|
| **AAEL009129** | cytochrome P450 | CYP6Z9 |
| **AAEL009630** | high-affinity cgmp-specific 3,5-cyclic phosphodiesterase | N/A |
| **AAEL009645** | unspecified product | N/A |
| **AAEL010075** | oxidoreductase | N/A |
| **AAEL010128** | leucine-rich immune protein (Long) | LRIM4 |
| **AAEL010366** | glucosyl/glucuronosyl transferases | N/A |
| **AAEL010712** | low-density lipoprotein receptor (ldl) | N/A |
| **AAEL011006** | guanylate kinase | N/A |
| **AAEL011133** | unspecified product | N/A |
| **AAEL011161** | unspecified product | N/A |
| **AAEL012110** | protease m1 zinc metalloprotease | N/A |
| **AAEL012409** | pantothenate kinase | N/A |
| **AAEL012446** | Inhibitor of Apoptosis (IAP) containing Baculoviral IAP Repeat(s) (BIR domains). | IAP6 |
| **AAEL012740** | ATPase subunit, putative | N/A |
| **AAEL013262** | unspecified product | N/A |
| **AAEL013347** | lethal(2)essential for life protein, l2efl | N/A |
| **AAEL013349** | lethal(2)essential for life protein, l2efl | N/A |
| **AAEL013431** | proline oxidase | N/A |
| **AAEL013484** | unspecified product | N/A |
| **AAEL013662** | anterior fat body protein | N/A |
| **AAEL013812** | unspecified product | N/A |
| **AAEL013885** | unspecified product | N/A |
| **AAEL014303** | neuroligin, | N/A |
| **AAEL014578** | ssm4 protein | N/A |
| **AAEL014619** | cytochrome P450 | CYP9J22 |
| **AAEL014863** | glycogenin | N/A |
| **AAEL014999** | unspecified product | N/A |
| **AAEL017514** | unspecified product | N/A |
| **AAEL018117** | unspecified product | N/A |
| **AAEL018219** | unspecified product | N/A |
| **AAEL018668** | ATP synthase F0 subunit 6 | ATP6 |
| **AAEL018685** | cytochrome b | CYTB |
| **AAEL019494** | unspecified product | N/A |
| **AAEL019495** | unspecified product | N/A |
| **AAEL019504** | unspecified product | N/A |
| **AAEL019639** | unspecified product | N/A |
| **AAEL019713** | unspecified product | N/A |
| **AAEL020524** | unspecified product | N/A |
| **AAEL020997** | pseudogene | N/A |
| **AAEL021035** | unspecified product | N/A |
| **AAEL021471** | unspecified product | N/A |
| **AAEL021762** | unspecified product | N/A |
| **AAEL021861** | unspecified product | N/A |
| **AAEL022059** | pseudogene | N/A |

| | | |
|---|---|---|
| **AAEL023634** | unspecified product | N/A |
| **AAEL023799** | pseudogene | N/A |
| **AAEL024512** | pseudogene | N/A |
| **AAEL025488** | unspecified product | N/A |
| **AAEL027008** | unspecified product | N/A |
| **AAEL027243** | pseudogene | N/A |
| **AAEL027593** | unspecified product | N/A |
| **AAEL027694** | unspecified product | N/A |
| **AAEL028635** | unspecified product | N/A |
| **AAEL029056** | unspecified product | N/A |

| Table S4. Genes differentially expressed in ZIKV-exposed COL.*w*Mel midguts 7dpf | | |
|---|---|---|
| **Gene ID** | **Product Description** | **Gene Name or Symbol** |
| **AAEL000080** | phosphoenolpyruvate carboxykinase | N/A |
| **AAEL000128** | P130 | N/A |
| **AAEL000271** | gamma-glutamyl hydrolase | N/A |
| **AAEL000294** | unspecified product | N/A |
| **AAEL000323** | cysteine-rich venom protein, putative | N/A |
| **AAEL000416** | FHA domain-containing protein [Source:UniProtKB/TrEMBL;Acc:A0A1S4EVV1] | N/A |
| **AAEL000488** | unspecified product | N/A |
| **AAEL000512** | Dynein heavy chain [Source:UniProtKB/TrEMBL;Acc:A0A1S4EW29] | N/A |
| **AAEL000566** | unspecified product | N/A |
| **AAEL000636** | unspecified product | N/A |
| **AAEL000713** | reticulon/nogo | N/A |
| **AAEL000757** | anterior fat body protein | N/A |
| **AAEL000859** | unspecified product | N/A |
| **AAEL000898** | unspecified product | N/A |
| **AAEL000902** | sugar transporter | N/A |
| **AAEL000905** | unspecified product | N/A |
| **AAEL000923** | unspecified product | N/A |
| **AAEL001062** | unspecified product | N/A |
| **AAEL001209** | sodium-dependent phosphate transporter | N/A |
| **AAEL001232** | tubulointerstitial nephritis antigen | N/A |
| **AAEL001254** | unspecified product | N/A |
| **AAEL001421** | high density lipoprotien binding protein / vigilin | N/A |
| **AAEL001434** | coronin | N/A |
| **AAEL001511** | unspecified product | N/A |
| **AAEL001532** | FAD NAD binding oxidoreductases | N/A |
| **AAEL001580** | otefin, putative | N/A |
| **AAEL001632** | multicopper oxidase | N/A |
| **AAEL001650** | ML domain-containing protein [Source:UniProtKB/TrEMBL;Acc:A0A1S4EZF1] | N/A |
| **AAEL001667** | multicopper oxidase | N/A |
| **AAEL001749** | ventrhoid transmembrane protein, putative | N/A |
| **AAEL001816** | glucosyl/glucuronosyl transferases | N/A |
| **AAEL001818** | unspecified product | N/A |
| **AAEL001837** | Lipase [Source:UniProtKB/TrEMBL;Acc:A0A1S4EZX5] | N/A |

| | | |
|---|---|---|
| **AAEL001900** | lactosylceramide 4-alpha-galactosyltransferase (alpha-1,4-galactosyltransferase) | N/A |
| **AAEL001905** | unspecified product | N/A |
| **AAEL001919** | protein tyrosine phosphatase, non-receptor type nt1 | N/A |
| **AAEL001986** | kinesin-like protein KIF1B | N/A |
| **AAEL002036** | unspecified product | N/A |
| **AAEL002080** | septin interacting protein, putative | N/A |
| **AAEL002102** | unspecified product | N/A |
| **AAEL002109** | unspecified product | N/A |
| **AAEL002130** | ecdysone inducible protein L2, putative | N/A |
| **AAEL002176** | inosine-uridine preferring nucleoside hydrolase | N/A |
| **AAEL002199** | unspecified product | N/A |
| **AAEL002235** | unspecified product | N/A |
| **AAEL002261** | GTP cyclohydrolase i | N/A |
| **AAEL002309** | Thioredoxin Peroxidase. | TPX4 |
| **AAEL002467** | unspecified product | N/A |
| **AAEL002557** | cationic amino acid transporter | N/A |
| **AAEL002623** | unspecified product | N/A |
| **AAEL002652** | unspecified product | N/A |
| **AAEL002661** | Matrix metalloproteinase [Source:UniProtKB/TrEMBL;Acc:A0A1S4F2E2] | N/A |
| **AAEL002671** | unspecified product | N/A |
| **AAEL002690** | beat protein | N/A |
| **AAEL002714** | kinesin-like protein KIF23 (mitotic kinesin-like protein 1) | N/A |
| **AAEL002757** | unspecified product | N/A |
| **AAEL002796** | l-asparaginase i | N/A |
| **AAEL002848** | tubulin beta chain | N/A |
| **AAEL002854** | F-box domain-containing protein [Source:UniProtKB/TrEMBL;Acc:A0A1S4F302] | N/A |
| **AAEL002886** | thioredoxin reductase | N/A |
| **AAEL002919** | unspecified product | N/A |
| **AAEL002921** | unspecified product | N/A |
| **AAEL002978** | leucyl aminopeptidase, putative | N/A |
| **AAEL003051** | unspecified product | N/A |
| **AAEL003063** | Semaphorin [Source:UniProtKB/TrEMBL;Acc:A0A1S4F3R5] | N/A |
| **AAEL003294** | fibrinogen and fibronectin | N/A |
| **AAEL003317** | alkaline phosphatase | N/A |
| **AAEL003589** | transcription factor, putative | N/A |
| **AAEL003619** | sodium/chloride dependent amino acid transporter | N/A |

| | | |
|---|---|---|
| **AAEL003681** | unspecified product | N/A |
| **AAEL003712** | C-Type Lysozyme (Lys-E). | LYSC10 |
| **AAEL003950** | helicase | N/A |
| **AAEL003951** | unspecified product | N/A |
| **AAEL004090** | unspecified product | N/A |
| **AAEL004092** | deoxyribonuclease I, putative | N/A |
| **AAEL004126** | sterol desaturase | N/A |
| **AAEL004206** | unspecified product | N/A |
| **AAEL004212** | unspecified product | N/A |
| **AAEL004302** | unspecified product | N/A |
| **AAEL004310** | p15-2a protein, putative | N/A |
| **AAEL004319** | epidermal growth factor receptor | N/A |
| **AAEL004386** | chorion peroxidase | pxt |
| **AAEL004392** | IAP-antagonist Michelob_x-like Protein | IMP |
| **AAEL004520** | cAMP/cgmp cyclic nucleotide phosphodiesterase | N/A |
| **AAEL004710** | spingomyelin synthetase | N/A |
| **AAEL004729** | unspecified product | N/A |
| **AAEL004868** | hemomucin | N/A |
| **AAEL004870** | cytochrome P450 | CYP18A1 |
| **AAEL004964** | unspecified product | N/A |
| **AAEL004981** | cation-transporting ATPase | N/A |
| **AAEL005008** | aquaporin | N/A |
| **AAEL005071** | GTP binding protein [Source:UniProtKB/TrEMBL;Acc:A0A1S4F9G8] | N/A |
| **AAEL005255** | PAR-domain protein 1 | PDP1 |
| **AAEL005342** | unspecified product | N/A |
| **AAEL005347** | unspecified product | N/A |
| **AAEL005417** | annexin x | N/A |
| **AAEL005428** | unspecified product | N/A |
| **AAEL005432** | unspecified product | N/A |
| **AAEL005455** | CTP synthase [Source:UniProtKB/TrEMBL;Acc:Q17A05] | CTPsyn |
| **AAEL005503** | unspecified product | N/A |
| **AAEL005666** | matrix metalloproteinase | N/A |
| **AAEL005701** | retinaldehyde binding protein | N/A |
| **AAEL005704** | unspecified product | N/A |
| **AAEL005791** | unspecified product | N/A |
| **AAEL005839** | uridine phosphorylase | N/A |
| **AAEL005977** | chondroitin 4-sulfotransferase | N/A |

| | | |
|---|---|---|
| **AAEL005992** | adam (a disintegrin and metalloprotease) | N/A |
| **AAEL006028** | unspecified product | N/A |
| **AAEL006034** | Vanin-like protein 1 precursor, putative | N/A |
| **AAEL006054** | peptidyl-prolyl cis-trans isomerase (cyclophilin) | N/A |
| **AAEL006171** | n-myc downstream regulated | N/A |
| **AAEL006216** | unspecified product | N/A |
| **AAEL006277** | unspecified product | N/A |
| **AAEL006321** | 1-acylglycerol-3-phosphate acyltransferase [Source:UniProtKB/TrEMBL;Acc:Q176M5] | N/A |
| **AAEL006355** | Class C Scavenger Receptor (Sushi/SCR/CCP MAM and Somatomedin B domains). | SCRC1 |
| **AAEL006361** | Class C Scavenger Receptor (Sushi/SCR/CCP MAM and Somatomedin B domains). | SCRC2 |
| **AAEL006449** | ser/thr protein kinase-lyk4 | N/A |
| **AAEL006480** | unspecified product | N/A |
| **AAEL006518** | cytidine deaminase, putative | N/A |
| **AAEL006663** | ANK_REP_REGION domain-containing protein [Source:UniProtKB/TrEMBL;Acc:A0A1S4FEA3] | N/A |
| **AAEL006686** | unspecified product | N/A |
| **AAEL006708** | hedgehog | N/A |
| **AAEL006723** | unspecified product | N/A |
| **AAEL006809** | voltage-gated ion channel | N/A |
| **AAEL006902** | serine-type enodpeptidase, | N/A |
| **AAEL006921** | calmodulin | N/A |
| **AAEL006978** | protein-glutamine gamma-glutamyltransferase | N/A |
| **AAEL007004** | GPCR Bride of Sevenless Family | GPRBOS1 |
| **AAEL007018** | udp-glucose 4-epimerase | N/A |
| **AAEL007030** | ceramidase | N/A |
| **AAEL007097** | 4-nitrophenylphosphatase [Source:UniProtKB/TrEMBL;Acc:Q0IF18] | N/A |
| **AAEL007120** | lim homeobox protein | N/A |
| **AAEL007191** | amino acid transporter | N/A |
| **AAEL007208** | unspecified product | N/A |
| **AAEL007238** | DUF3421 domain-containing protein [Source:UniProtKB/TrEMBL;Acc:A0A1S4FG14] | N/A |
| **AAEL007258** | unspecified product | N/A |
| **AAEL007271** | basic helix-loop-helix zip transcription factor | N/A |
| **AAEL007299** | Cadherin, putative [Source:UniProtKB/TrEMBL;Acc:A0A1S4FG21] | N/A |
| **AAEL007344** | LITAF domain-containing protein [Source:UniProtKB/TrEMBL;Acc:A0A1S4FG64] | N/A |
| **AAEL007547** | chloride channel protein | N/A |

| | | |
|---|---|---|
| **AAEL007560** | core 1 udp-galactose:n-acetylgalactosamine-alpha-r beta 1,3-galactosyltransferase | N/A |
| **AAEL007657** | low-density lipoprotein receptor (ldl) | N/A |
| **AAEL007765** | Serine Protease Inhibitor (serpin) likely cleavage at K/R. Transcript A. | SRPN10 |
| **AAEL007778** | leucine-rich transmembrane protein | N/A |
| **AAEL007872** | unspecified product | N/A |
| **AAEL007880** | ornithine decarboxylase | N/A |
| **AAEL007942** | fibrinogen and fibronectin | N/A |
| **AAEL007993** | Clip-Domain Serine Protease family B. | CLIPB27 |
| **AAEL008024** | unspecified product | N/A |
| **AAEL008027** | unspecified product | N/A |
| **AAEL008028** | monocarboxylate transporter | N/A |
| **AAEL008097** | trypsin-eta, putative | N/A |
| **AAEL008141** | period circadian protein | PER |
| **AAEL008267** | GPCR Neurokinin/Tachykinin Family | GPRNPR5 |
| **AAEL008306** | mitogen activated protein kinase kinase kinase 5, mapkkk5, mekk5 | N/A |
| **AAEL008346** | achaete-scute complex protein T3, putative | N/A |
| **AAEL008467** | cysteine synthase | N/A |
| **AAEL008468** | cysteine synthase | N/A |
| **AAEL008473** | cysteine-rich venom protein, putative | N/A |
| **AAEL008511** | unspecified product | N/A |
| **AAEL008547** | unspecified product | N/A |
| **AAEL008622** | jnk | N/A |
| **AAEL008655** | GPCR Vasopressin Family | GPRVPR2 |
| **AAEL008658** | leucine-rich immune protein (TM) | LRIM16 |
| **AAEL008760** | unspecified product | N/A |
| **AAEL008767** | serine protease | N/A |
| **AAEL008782** | serine-type enodpeptidase, | N/A |
| **AAEL008829** | unspecified product | N/A |
| **AAEL008832** | forkhead box protein (AaegFOXN1) | N/A |
| **AAEL008843** | unspecified product | N/A |
| **AAEL008910** | unspecified product | N/A |
| **AAEL008916** | unspecified product | N/A |
| **AAEL008921** | myosin regulatory light chain 2 smooth muscle | N/A |
| **AAEL008953** | unspecified product | N/A |
| **AAEL009070** | unspecified product | N/A |
| **AAEL009114** | unspecified product | N/A |
| **AAEL009185** | arginine or creatine kinase | N/A |

| | | |
|---|---|---|
| **AAEL009249** | Coronin [Source:UniProtKB/TrEMBL;Acc:A0A1S4FM51] | N/A |
| **AAEL009317** | rab11 | N/A |
| **AAEL009333** | unspecified product | N/A |
| **AAEL009371** | unspecified product | N/A |
| **AAEL009556** | Niemann-Pick Type C-2, putative | N/A |
| **AAEL009645** | unspecified product | N/A |
| **AAEL009681** | Putative rhomboid family [Source:UniProtKB/TrEMBL;Acc:A0A0P6IZ43] | N/A |
| **AAEL009762** | cytochrome P450 | CYP307A1 |
| **AAEL009813** | glutamate receptor 7 (ampa) | N/A |
| **AAEL009842** | Galectin [Source:UniProtKB/TrEMBL;Acc:Q16UP1] | GALE12 |
| **AAEL009850** | Galectin [Source:UniProtKB/TrEMBL;Acc:Q16UP0] | GALE14 |
| **AAEL009987** | unspecified product | N/A |
| **AAEL010050** | unspecified product | N/A |
| **AAEL010075** | oxidoreductase | N/A |
| **AAEL010084** | unspecified product | N/A |
| **AAEL010094** | cyclin b | N/A |
| **AAEL010145** | sodium/potassium-dependent ATPase beta-2 subunit | N/A |
| **AAEL010264** | unspecified product | N/A |
| **AAEL010270** | unspecified product | N/A |
| **AAEL010375** | unspecified product | N/A |
| **AAEL010477** | unspecified product | N/A |
| **AAEL010483** | oxysterol-binding protein related protein (ORP8) | ORP8 |
| **AAEL010650** | sodium/solute symporter | N/A |
| **AAEL010661** | phospholipid scramblase 1, | N/A |
| **AAEL010678** | unspecified product | N/A |
| **AAEL010738** | sodium bicarbonate cotransporter | N/A |
| **AAEL010776** | carboxypeptidase | N/A |
| **AAEL010782** | carboxypeptidase | N/A |
| **AAEL010840** | unspecified product | N/A |
| **AAEL010932** | RNAse h | N/A |
| **AAEL010956** | unspecified product | N/A |
| **AAEL011009** | fibrinogen and fibronectin | N/A |
| **AAEL011203** | unspecified product | N/A |
| **AAEL011264** | phosphatidylethanolamine-binding protein | N/A |
| **AAEL011424** | Histone H3 [Source:UniProtKB/TrEMBL;Acc:A0A1S4FTJ0] | N/A |
| **AAEL011510** | multiple inositol polyphosphate phosphatase | N/A |
| **AAEL011598** | Gustatory receptor [Source:UniProtKB/TrEMBL;Acc:A0A1S4FU22] | N/A |

| | | |
|---|---|---|
| **AAEL011648** | cyclin d | N/A |
| **AAEL011650** | coatomer, gamma-subunit, putative | N/A |
| **AAEL011653** | thyroid hormone receptor interactor | N/A |
| **AAEL011901** | 1-acyl-sn-glycerol-3-phosphate acyltransferase | N/A |
| **AAEL011937** | glucosyl/glucuronosyl transferases | N/A |
| **AAEL012003** | Galectin [Source:UniProtKB/TrEMBL;Acc:Q17AG2] | GALE6B |
| **AAEL012014** | L-lactate dehydrogenase [Source:UniProtKB/TrEMBL;Acc:Q16ND1] | N/A |
| **AAEL012052** | unspecified product | N/A |
| **AAEL012062** | Na+/K+ ATPase alpha subunit | N/A |
| **AAEL012349** | lipase 1 precursor | N/A |
| **AAEL012390** | unspecified product | N/A |
| **AAEL012410** | eukaryotic translation initiation factor 2C | AGO1b |
| **AAEL012499** | Histone H2A [Source:UniProtKB/TrEMBL;Acc:Q16LW9] | N/A |
| **AAEL012514** | translation initiation factor 2b, delta subunit | N/A |
| **AAEL012522** | Sodium-dependent phosphate transporter [Source:UniProtKB/TrEMBL;Acc:A0A1S4FWV3] | N/A |
| **AAEL012545** | Proliferating cell nuclear antigen [Source:UniProtKB/TrEMBL;Acc:Q4PKD7] | N/A |
| **AAEL012629** | deoxyuridine 5'-triphosphate nucleotidohydrolase | N/A |
| **AAEL012859** | unspecified product | N/A |
| **AAEL012960** | importin alpha | N/A |
| **AAEL013111** | glutamate transporter | N/A |
| **AAEL013262** | unspecified product | N/A |
| **AAEL013276** | acid phosphatase | N/A |
| **AAEL013304** | unspecified product | N/A |
| **AAEL013309** | high-affinity copper uptake protein | N/A |
| **AAEL013345** | alphaA-crystallin, putative | N/A |
| **AAEL013346** | lethal(2)essential for life protein, l2efl | N/A |
| **AAEL013348** | lethal(2)essential for life protein, l2efl | N/A |
| **AAEL013349** | lethal(2)essential for life protein, l2efl | N/A |
| **AAEL013350** | heat shock protein 26kD, putative | N/A |
| **AAEL013351** | lethal(2)essential for life protein, l2efl | N/A |
| **AAEL013352** | lethal(2)essential for life protein, l2efl | N/A |
| **AAEL013692** | PIWI | PIWI3 |
| **AAEL013713** | trypsin | N/A |
| **AAEL013780** | unspecified product | N/A |
| **AAEL013808** | fascin | N/A |
| **AAEL013875** | tetraspanin, putative | N/A |
| **AAEL014019** | cytochrome P450 | CYP4J16 |

| | | |
|---|---|---|
| **AAEL014226** | unspecified product | N/A |
| **AAEL014251** | Inhibitor of Apoptosis (IAP) containing Baculoviral IAP Repeat(s) (BIR domains). | IAP5 |
| **AAEL014348** | caspase (short) | CASPS8 |
| **AAEL014363** | unspecified product | N/A |
| **AAEL014408** | m-phase inducer phosphatase(cdc25) | N/A |
| **AAEL014439** | juvenile hormone-inducible protein, putative | N/A |
| **AAEL014454** | unspecified product | N/A |
| **AAEL014541** | maltose phosphorylase | N/A |
| **AAEL014567** | oviductin | N/A |
| **AAEL014945** | unspecified product | N/A |
| **AAEL014981** | unspecified product | N/A |
| **AAEL016975** | unspecified product | N/A |
| **AAEL017331** | unspecified product | N/A |
| **AAEL017553** | Carboxy/choline esterase Alpha Esterase | CCEAE2B |
| **AAEL018039** | unspecified product | N/A |
| **AAEL018118** | unspecified product | N/A |
| **AAEL018120** | Ribosomal protein S6 kinase [Source:UniProtKB/TrEMBL;Acc:Q535V4] | N/A |
| **AAEL019504** | unspecified product | N/A |
| **AAEL019536** | unspecified product | N/A |
| **AAEL019604** | unspecified product | N/A |
| **AAEL019610** | unspecified product | N/A |
| **AAEL019623** | unspecified product | N/A |
| **AAEL019677** | unspecified product | N/A |
| **AAEL019681** | unspecified product | N/A |
| **AAEL019700** | unspecified product | N/A |
| **AAEL019712** | unspecified product | N/A |
| **AAEL019722** | unspecified product | N/A |
| **AAEL019728** | suppressor of cytokine signaling | SOCS |
| **AAEL019785** | unspecified product | N/A |
| **AAEL019793** | unspecified product | N/A |
| **AAEL019902** | unspecified product | N/A |
| **AAEL019903** | unspecified product | N/A |
| **AAEL019940** | unspecified product | N/A |
| **AAEL019941** | unspecified product | N/A |
| **AAEL020654** | unspecified product | N/A |
| **AAEL020729** | unspecified product | N/A |
| **AAEL020800** | unspecified product | N/A |

| | | |
|---|---|---|
| **AAEL021278** | unspecified product | N/A |
| **AAEL021318** | unspecified product | N/A |
| **AAEL021321** | unspecified product | N/A |
| **AAEL021899** | unspecified product | N/A |
| **AAEL021925** | unspecified product | N/A |
| **AAEL021931** | unspecified product | N/A |
| **AAEL022048** | unspecified product | N/A |
| **AAEL022059** | pseudogene | N/A |
| **AAEL022167** | unspecified product | N/A |
| **AAEL022628** | unspecified product | N/A |
| **AAEL022659** | unspecified product | N/A |
| **AAEL023231** | unspecified product | N/A |
| **AAEL023478** | unspecified product | N/A |
| **AAEL023490** | unspecified product | N/A |
| **AAEL023644** | unspecified product | N/A |
| **AAEL023746** | unspecified product | N/A |
| **AAEL024003** | unspecified product | N/A |
| **AAEL024222** | unspecified product | N/A |
| **AAEL024370** | unspecified product | N/A |
| **AAEL024512** | pseudogene | N/A |
| **AAEL024520** | unspecified product | N/A |
| **AAEL024558** | unspecified product | N/A |
| **AAEL024675** | unspecified product | N/A |
| **AAEL024880** | unspecified product | N/A |
| **AAEL024913** | unspecified product | N/A |
| **AAEL024940** | unspecified product | N/A |
| **AAEL025091** | unspecified product | N/A |
| **AAEL025226** | unspecified product | N/A |
| **AAEL025432** | Cytidine deaminase [Source:UniProtKB/TrEMBL;Acc:Q0IF52] | N/A |
| **AAEL025530** | unspecified product | N/A |
| **AAEL025658** | unspecified product | N/A |
| **AAEL025718** | unspecified product | N/A |
| **AAEL025818** | unspecified product | N/A |
| **AAEL025839** | unspecified product | N/A |
| **AAEL026025** | unspecified product | N/A |
| **AAEL026215** | unspecified product | N/A |
| **AAEL026343** | unspecified product | N/A |

| | | |
|---|---|---|
| **AAEL026466** | unspecified product | N/A |
| **AAEL026744** | unspecified product | N/A |
| **AAEL026751** | unspecified product | N/A |
| **AAEL026981** | unspecified product | N/A |
| **AAEL027019** | unspecified product | N/A |
| **AAEL027398** | unspecified product | N/A |
| **AAEL027593** | unspecified product | N/A |
| **AAEL027610** | unspecified product | N/A |
| **AAEL027937** | unspecified product | N/A |
| **AAEL028005** | unspecified product | N/A |
| **AAEL028021** | unspecified product | N/A |
| **AAEL028048** | unspecified product | N/A |
| **AAEL028088** | unspecified product | N/A |
| **AAEL028635** | unspecified product | N/A |
| **AAEL029041** | unspecified product | N/A |
| **AAEL029044** | cecropin | CECE |
| **AAEL029046** | cecropin | CECD |
| **AAEL029082** | unspecified product | N/A |
| **AAEL029104** | unspecified product | N/A |
| **AAEL029107** | unspecified product | N/A |
